# The Physiological Landscape and Specificity of Antibody Repertoires

**DOI:** 10.1101/2021.09.15.460420

**Authors:** Lucia Csepregi, Kenneth B. Hoehn, Daniel Neumeier, Joseph M. Taft, Simon Friedensohn, Cédric R. Weber, Arkadij Kummer, Fabian Sesterhenn, Bruno E. Correia, Sai T. Reddy

## Abstract

Diverse antibody repertoires spanning multiple lymphoid organs (e.g., bone marrow, spleen, lymph nodes) form the foundation of protective humoral immunity. Changes in their composition across lymphoid organs are a consequence of B-cell selection and migration events leading to a highly dynamic and unique physiological landscape of antibody repertoires upon antigenic challenge (e.g., vaccination). However, to what extent B cells encoding identical or similar antibody sequences (clones) are distributed across multiple lymphoid organs and how this is shaped by the strength of a humoral response, remains largely unexplored. Here, we performed an in-depth systems analysis of antibody repertoires across multiple distinct lymphoid organs of immunized mice, and discovered that organ-specific antibody repertoire features (e.g., germline V-gene usage and clonal expansion profiles) equilibrated upon a strong humoral response (multiple immunizations and high serum titers). This resulted in a surprisingly high degree of repertoire consolidation, characterized by highly connected and overlapping B-cell clones across multiple lymphoid organs. Finally, we revealed distinct physiological axes indicating clonal migrations and showed that antibody repertoire consolidation directly correlated with antigen-specificity. Our study uncovered how a strong humoral response resulted in a more uniform but redundant physiological landscape of antibody repertoires, indicating that increases in antibody serum titers were a result of synergistic contributions from antigen-specific B-cell clones distributed across multiple lymphoid organs. Our findings provide valuable insights for the assessment and design of vaccine strategies.

## Introduction

B cells exert their protective functions via their B-cell receptors (BCR) [and secreted version: antibodies], which are capable of recognizing a plethora of antigenic epitopes and structures. Hematopoietic precursors differentiate into naive B cells in the bone marrow, which then migrate to secondary lymphoid organs such as the spleen, lymph nodes and mucosa-associated lymphoid tissue, which are distributed throughout the body and are interconnected through the circulatory and lymphatic systems. Naive B cells, which are continuously circulating and screening for antigens, can become activated upon an encounter with a cognate antigen (BCR binding to antigen), which results in differentiation into memory B cells and plasma cell subsets that eventually migrate to supportive niches for their long-term survival (*1, 2*). Thus, the composition of antigen-specific B cells throughout the body is highly dynamic on a molecular, cellular and spatio-temporal level, resulting in a highly diverse and unique physiological landscape of antibody repertoires (*3*).

In order to assess antigen-induced B-cell responses, measurements of serum antibody titers can be performed, which are mainly reflective of circulating antibodies [immunoglobulins (Ig)] secreted from long-lived plasma cells that reside in the bone marrow, spleen and lymph nodes (*1, 4, 5*). However, it is still unknown how the strength of humoral immunity, which is correlated with serum antibody titers (e.g., IgG), is reflected in the physiological composition of antibody repertoires present across multiple lymphoid organs.

Thus far, only a few systematic studies have profiled antibody repertoires across multiple organs within the same individuals. In humans, due to limited organ accessibility, most studies perform antibody repertoire analysis on B cells derived from peripheral blood, with a few notable exceptions analyzing more than two organs (*6–9*). For example, Meng et al. constructed an atlas of B-cell clonal distribution uncovering distinct evolution of clonal lineages in human tissues such as blood-rich compartments versus the gastrointestinal tract (*7*). Yang et al. reported shared antibody repertoires across human tissues and inferred antigen-specificity based on similarity to previously reported pathogen-specific antibody sequences (*8*). In the context of mice, Mathew et al. analyzed antigen-specific B cells in the draining lymph nodes, spleen and lungs after influenza infection, which revealed distinct organ-specific antibody repertoire features (*10*).

However, similar to other studies across tissues in mice, they performed small-scale sequencing on specific B-cell subsets (*11–13*). Therefore, it remains an open question how the degree of clonal overlap across physiological compartments relates on repertoire-scale and whether each lymphoid organ represents an exclusive or redundant (class-switched IgG) antibody repertoire upon antigenic challenge.

Here, we map the physiological landscape of antibody repertoires in mice following immunization. We used a systems-based approach that included deep and single-cell sequencing, bioinformatic and statistical analysis and high-throughput antibody specificity screening to comprehensively profile antibody repertoires from six distinct lymphoid organs. Our analysis uncovered that strong humoral responses (high antibody titers due to repeated immunizations) led to the physiological consolidation of antibody repertoires (a high fraction of clones shared across multiple lymphoid organs), which was directly correlated with antigen-binding specificity. The discovery that strong humoral immunity corresponds to a more uniform and homogenous physiological antibody repertoire provides important insights for the assessment and design of vaccine strategies.

## Results

### Systems analysis of the physiological landscape of antibody repertoires

To profile the physiological landscape of antibody repertoires, we performed a comprehensive systems analysis that included murine models of immunization, antibody repertoire sequencing, bioinformatic repertoire analysis and high-throughput antibody specificity screening (**Fig. 1A, fig. 5, A** and **G)**. The study was designed around two cohorts of mice that either received a single immunization (cohort-1x, consisting of mice 1x-A, 1x-B and 1x-C) or multiple booster immunizations (cohort-3x, mice 3x-D, 3x-E and 3x-F). Mice were subcutaneously injected at the left lateral flank with viral protein antigen [(pre-fusion glycoprotein of respiratory syncytial virus (RSV-F)] in poly(I:C) adjuvant, with cohort-3x mice receiving three injections at 21 day intervals (**Fig. 1A**), as it is well-established that multiple booster immunization schemes induce strong humoral immune responses (*4, 14*). For both cohorts, mice were sacrificed ten days after the final immunization, followed by isolation of six lymphoid organs: bone marrow, spleen and left and right axillary and inguinal lymph nodes. Antigen-binding titers from serum confirmed that cohort-3x mice indeed had a strong antibody response (average endpoint titer > 1 / 1 x 10^6^) relative to the weaker response of cohort-1x mice (average endpoint titer ∼ 1 / 3 x 10^4^) (**Fig. S1A**).

**Fig. 1.**
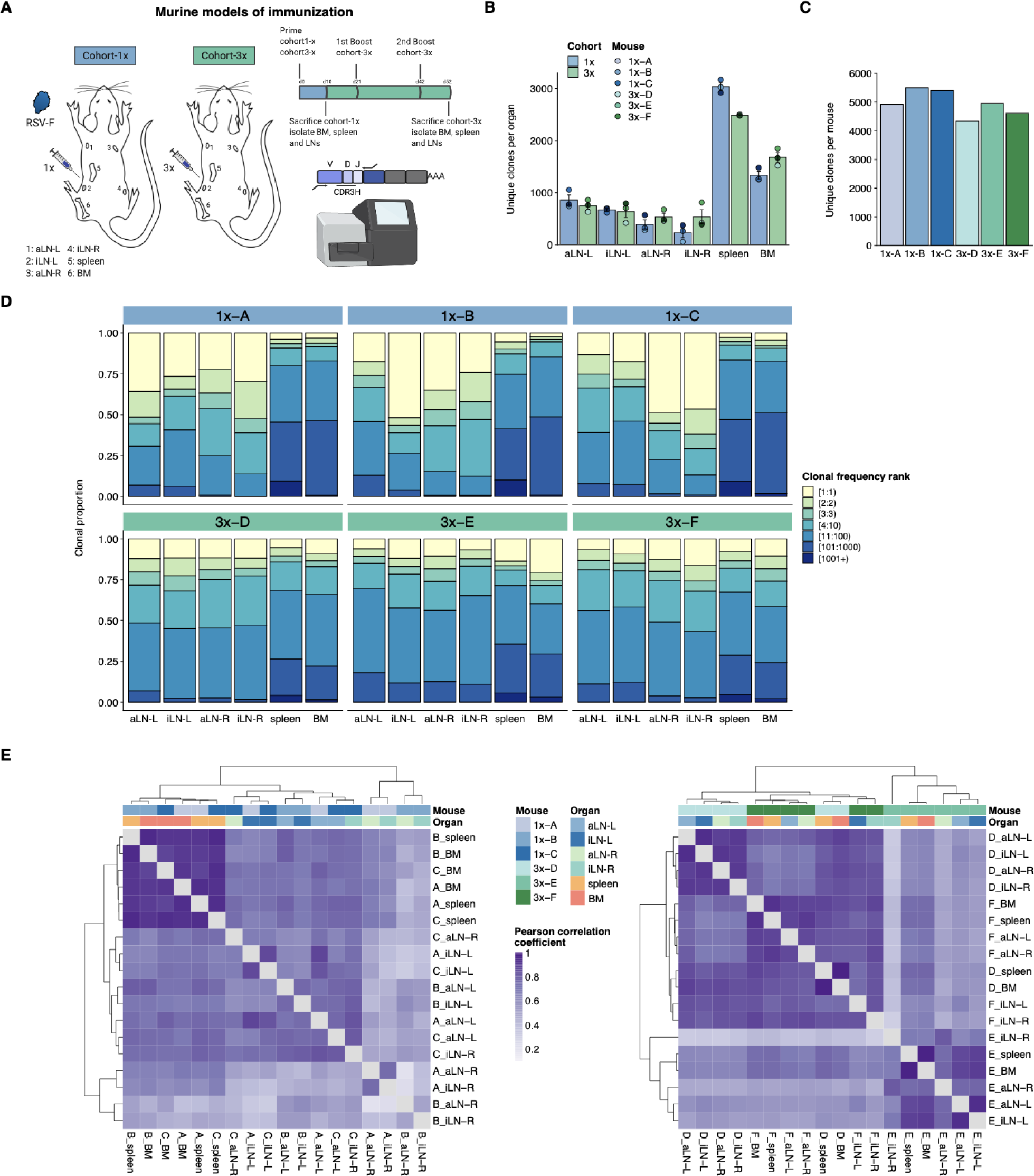
Profiling antibody repertoire diversity and clonal expansion across multiple lymphoid organs. (**A**) Overview of mouse immunizations with RSV-F antigen: mice (n=3 per cohort) were subjected to single (cohort-1x) or multiple (cohort-3x) immunizations (subcutaneous injections into the left lateral flank), followed by deep sequencing of antibody repertoires (VH IgG) derived from six different lymphoid organs [bone marrow (BM), spleen, left and right axillary lymph nodes (aLN-L, -R) and left and right inguinal lymph nodes (iLN-L, -R)]. (**B**) Shown is the average clonal diversity (mean ± SEM, n=3) based on the total number of unique clones within each organ per mouse (point) and cohort (bar). (**C**) Clonal diversity by quantification of total numbers of unique clones across all organs within each mouse. (**D**) Clonal expansion profiles with each bar representing an organ and each colored section displaying the repertoire space occupied by frequency ranked clones or clonal fractions within each organ. (**E**) Hierarchical clustering of Pearson correlation coefficients of germline V-gene usage counts between organs. Cohort-1x and cohort-3x were separately clustered (left and right). Each tile represents the pairwise Pearson correlation coefficient of the germline V-gene repertoires of two organs.

Deep sequencing of antibody repertoires has emerged as a powerful tool for assessing important immunological parameters such as clonal diversity, expansion and evolution (somatic hypermutation) and thus offers a quantitative approach to profile antibody repertoires from specific tissues (*15–18*). In order to reveal the physiological landscape of antibody repertoires following immunization, we performed targeted deep sequencing of six lymphoid organs in all three mice of both cohorts (total of 36 organs). Variable heavy (VH) chain IgG sequencing libraries were prepared according to our previously established protocol that incorporates unique molecular identifiers for error and bias correction (*19*). Focusing on the IgG-class-switched repertoire allowed for a pooled analysis of mainly plasmablasts and plasma cells due to their higher rate of antibody gene transcription. Deep sequencing (Illumina MiSeq, paired-end 2 x 300bp) and pre-processing resulted in an average of 6 x 10^5^ reads per organ and up to 4 x 10^5^ quality processed and merged reads per organ were used as input for the error-correction pipeline (see **Table S1**).

### Antibody repertoire analysis across lymphoid organs

First, we aimed to characterize common antibody repertoire features (e.g., clonal diversity, expansion and germline V-gene usage) in each lymphoid organ for both immunization cohorts. For this, we defined B-cell clones as antibody sequences possessing identical germline V- and J-genes and 100% CDRH3 amino acid (a.a.) identity and length. For investigation of groups of clonally related B-cell variants (here referred to as clonotypes), antibody sequences were clustered together in the same manner with the exception of using 90% CDRH3 a.a. identity (similar ratios between clones and clonotypes were observed when using 80% or 85% CDRH3 a.a. identity, see **Fig. S1B**).

Quantification of unique clones showed that lymph nodes, which are smaller in size, displayed less diverse antibody repertoires (55-1055 unique clones) than bone marrow (1252-1846 unique clones) and spleen (2475-3166 unique clones) (**Fig. 1B**, for diversity of unique clonotypes see **Table S2** and **Fig. S1C**). Comparing the diversity of clones per organ across both immunization cohorts showed that the left lymph nodes (side of immunization) displayed slightly higher diversities compared to the right lymph nodes. This difference was not visible in cohort-3x mice, which exhibited comparable diversities in lymph nodes from both sides. Moreover, whereas the bone marrow repertoires did not show cohort-dependent differences, spleen repertoires displayed reduced clonal diversity in cohort-3x mice. Pooling unique clones or clonotypes derived from all six lymphoid organs within each mouse revealed that cohort-3x mice had a reduced antibody repertoire diversity per mouse (**Fig. 1C** and **fig. S1D**). To account for differences in sequencing input across each organ, subsampling analysis was performed with varying sequence input into the error-correction pipeline, which confirmed the observed differences in diversity profiles across organs (**Fig. S1, E** and **F**).

Next, we assessed the distribution of clonal frequencies in all organ repertoires using Hill-based Evenness profiles (*20*), which revealed a high degree of clonal expansion in lymph nodes compared to spleen and bone marrow in cohort-1x mice. By contrast, expansion was uniformly polarized across all organs in cohort-3x mice (**Fig. S2A**). To further dissect what fraction of clones contributed to the observed repertoire polarization, we performed clonal expansion analysis by integrating clonal frequency information for specific proportions of clones present in each organ (**Fig. 1D**). Notably, both left and right lymph node repertoires of cohort-1x mice were dominated by only a few clones, which showed massive expansion. For instance, the three most frequent clones comprised 33% to 71% of the whole repertoire in all three mice. This stands in contrast to the spleen and bone marrow, in which the three most frequent clones occupied only up to 13% and 10%, respectively. Strikingly, compared to cohort-1x, expansion profiles in cohort-3x were more uniform across all organs with the three most expanded clones displaying frequencies ranging from 15% to 32% in lymph nodes and up to 19% and 29% in spleen and bone marrow, respectively.

Next, quantification of germline V-gene usage of unique clones revealed that clones from spleen and bone marrow contained more unique V-genes (99–115) compared to clones from lymph nodes (31–92), which is in line with the observed reduction of clonal diversity in lymph nodes of all mice. Hierarchical clustering of Pearson correlation coefficients for each mouse showed that spleen and bone marrow displayed the highest correlation in germline V-gene usage across both cohorts (**Fig. S2B**). In two out of three mice per cohort, the left lymph nodes as well as right lymph nodes clustered together, respectively, possibly reflecting a physiological axis upon immunization. Moreover, we observed cohort-specific differences of germline V-gene usage (**Fig. 1E**). In cohort-1x mice, the same organs of different mice clustered together, with the spleen and bone marrow displaying the highest similarity. By contrast, in cohort-3x mice, organs within the same mouse clustered together, reflecting a physiological consolidation of germline V-gene repertoires.

### Strong humoral responses result in highly connected and overlapping clones

In addition to clonal expansion, antigen immunization drives affinity maturation by somatic hypermutation, which results in production of clonal variants. To understand the network of clonal variants, we constructed sequence-similarity networks to elucidate clonal relationships (*21, 22*). Networks were generated where clones (nodes) within a clonotype were connected (edges) if they differed by one a.a. in CDRH3 (**Fig. 2,** legend). When assessing clonal connectivity by quantifying the number of edges per node, cohort-3x mice displayed an increased fraction of clones with higher connectivity compared to cohort-1x mice (**Fig. S3A**).

**Fig. 2.**
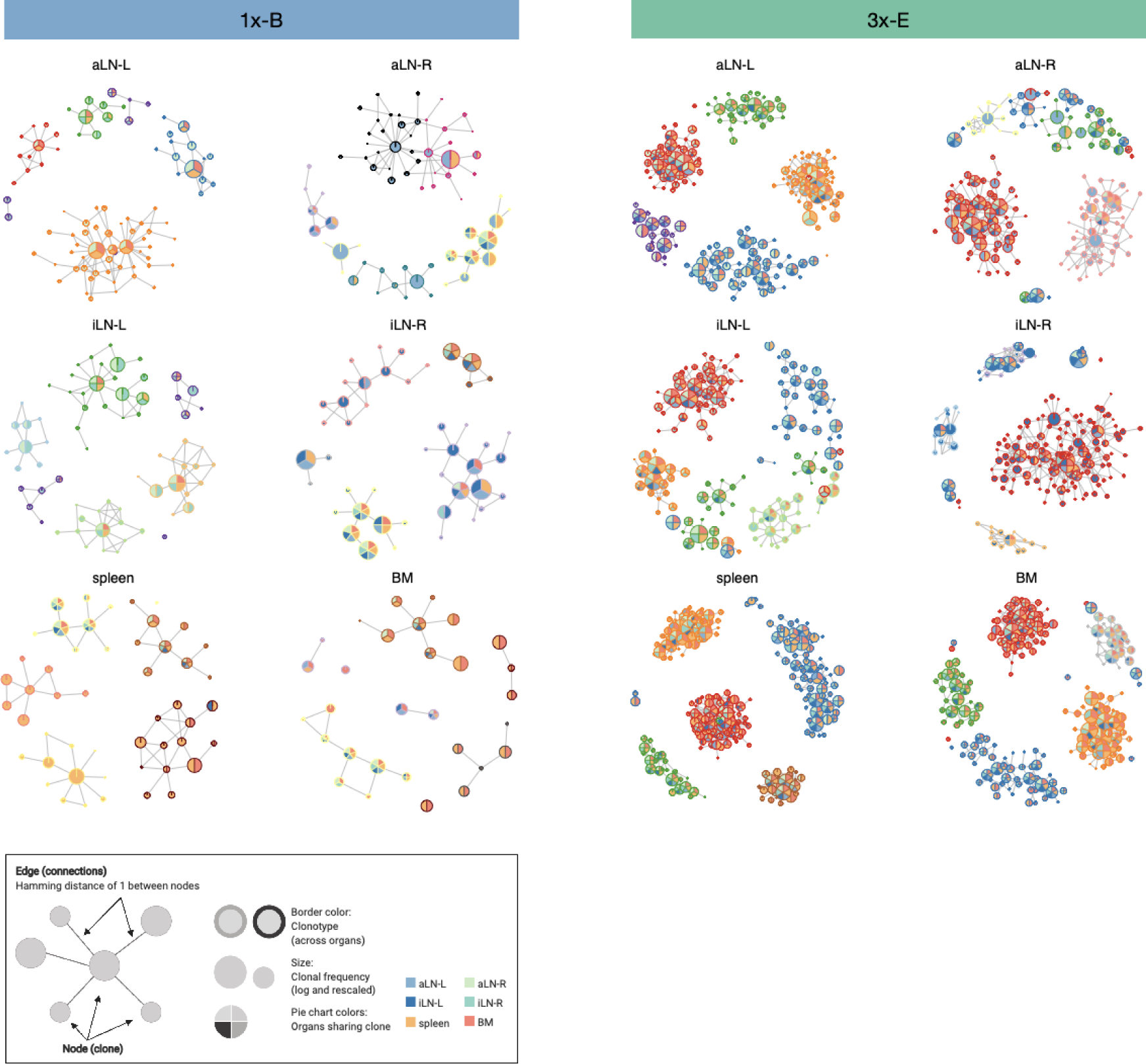
Sequence-similarity network analysis across lymphoid organs. Sequence-similarity networks of the five most diverse clonotypes from a representative mouse from the single (cohort-1x) or multiple (cohort-3x) immunization cohorts. Within a network, each node represents a unique clone that is connected by edges to clones with a Hamming distance of 1 between CDRH3 a.a. Clones within each clonotype display the same border color throughout all organs in each mouse. The size of each clone represents the clonal frequency (in log) within the corresponding organ. Pie charts within each node depict organs sharing the corresponding clone (see legend). See **Fig. S3C** for sequence similarity networks of the other cohort-1x and cohort-3x mice. BM: bone marrow, aLN-L, -R: left and right axillary lymph nodes, iLN-L, -R: left and right inguinal lymph nodes.

Network visualization of the five most diverse clonotypes per organ revealed that cohort-3x mice contained a substantially higher clonal connectivity and diversity across all organs, in particular for the spleen and bone marrow (**Fig. 2, Fig. S3, B** and **C**). These observations were in line with higher somatic hypermutation counts across all organs in cohort-3x mice (**Fig. S2C**). Notably, networks of the five most diverse clonotypes of both cohorts consisted of low and high frequency clones, with frequencies ranging from 5x10^-4^ % and 6x10^-4^ % to 22% and 21% within the respective organs in cohort-1x and cohort-3x mice, respectively. Moreover, within cohort-3x mice a substantial fraction of the five most diverse clonotypes were shared across all six organs (**Table S3**), and the majority of clones were present in all organs (**Fig. 2** and **fig. S3C**).

To get deeper insights into global repertoire similarities across all six organs within each mouse, we performed extensive clonal overlap analysis. Whereas in cohort-1x mice the majority of clones were found in a single organ (organ-exclusive), in cohort-3x mice clones were frequently shared across lymphoid organs, particularly the lymph nodes (**Fig. 3A**). Upon examining all unique clones, there was minimal overlap in cohort-1x mice, with only 1-3 clones being shared across all six organs (**Fig. 3B** and **fig. S4A**). In contrast, we observed extensive sharing across all lymphoid organs in cohort-3x mice, with 49-71 shared clones. Cohort-3x mice also showed a greater degree of repertoire similarity across all organs based on pairwise Jaccard correlation (clonal identity) and cosine similarity measurements (clonal identity and frequency) (**Fig. S4B, Fig. 3C** and **fig. S5**). This is consistent with the observation of higher similarities in germline V-gene usage across organs in cohort-3x mice. Physiological segregation of clones was observed with the left lymph nodes, right lymph nodes, as well as the spleen and bone marrow displaying increased repertoire similarities (based on germline V-gene usage and clonal overlap), compared to other organs across both cohorts. Of note, mouse 1x-C showed high pairwise cosine similarity between the right inguinal and axillary lymph nodes because both organs contained an identical clone (CDRH3: CARSLYGAFDYW) representing 47% and 49% of the repertoire, respectively. Focusing on the ten most frequent clones for each organ showed that shared clones also displayed similar frequencies across organs in cohort-3x and that increased overlap was also detected in low frequency clone fractions compared to cohort-1x (**Fig. S6**).

**Fig. 3.**
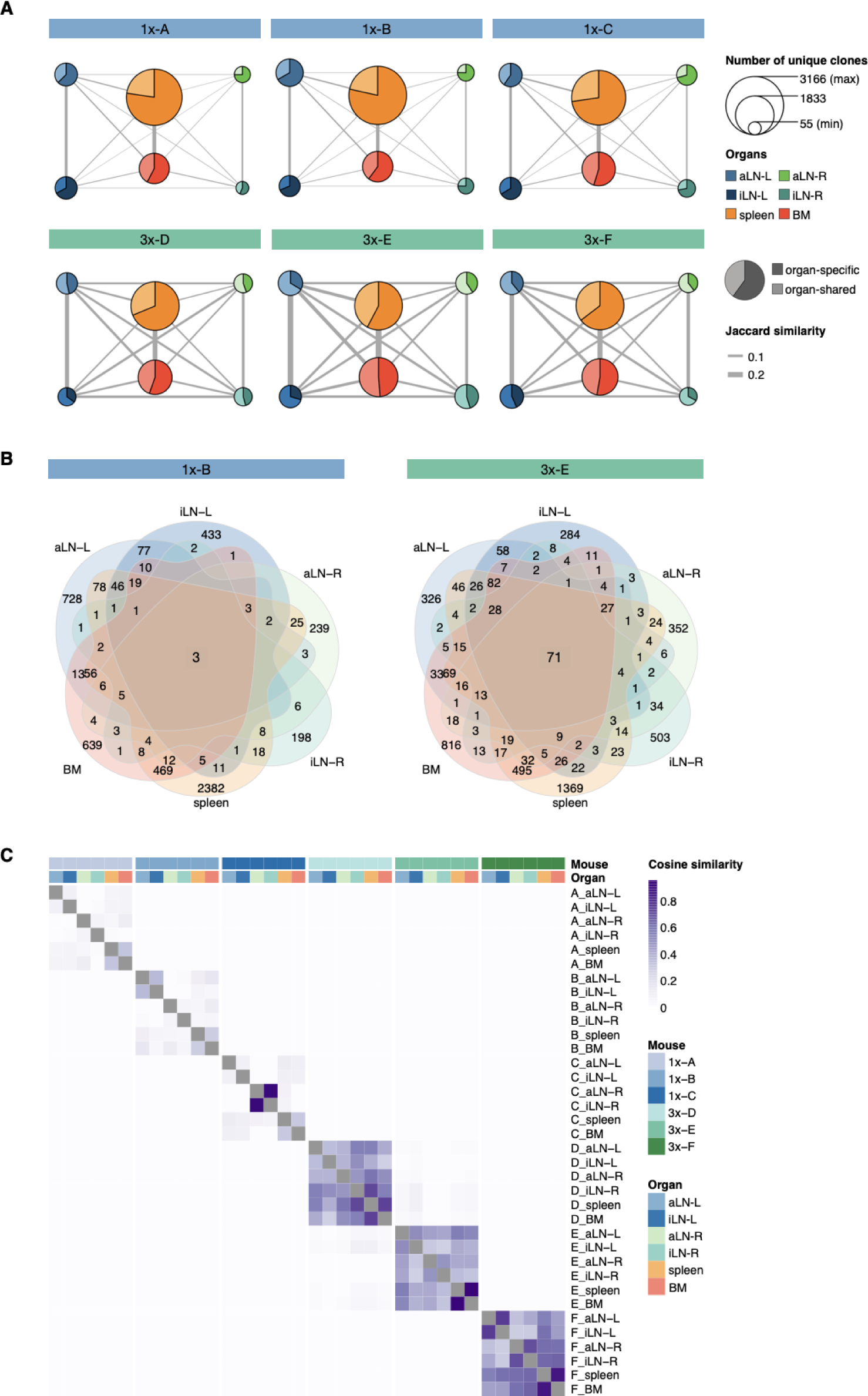
Strong humoral responses result in increased antibody repertoire similarities across multiple lymphoid organs. (**A**) Lymphoid organ repertoire similarity networks are based on clonal overlap across all six lymphoid organs per mouse. Each node represents an organ, with the size of the node being proportional to the number of unique clones within the organ. Width of edges between the nodes depict the pairwise repertoire similarity based on Jaccard similarity (see methods). The pie chart within each node represents the proportion of organ-specific and organ-shared clones for each organ. (**B**) Venn diagrams depicting numbers of shared unique clones from a representative mouse from the single (cohort-1x) or multiple (cohort-3x) immunization cohorts. See **Fig. S4A** for venn diagrams of the other cohort-1x and cohort-3x mice. (**C**) Heatmap representing pairwise cosine similarity indices based on clonal overlap including clonal frequency information across all samples. BM: bone marrow, aLN-L, -R: left and right axillary lymph nodes, iLN-L, -R: left and right inguinal lymph nodes.

### Phylogenetic analysis reveals physiological axes of clonal evolution

We next aimed to understand the evolution of B cells across all six lymphoid organs during weak and strong humoral responses. To trace the evolution of clones across multiple organs, we inferred lineage trees for each clonotype using *IgPhyML* (*23*). Lineage trees containing clonal sequences from multiple organs may represent clonotypes that have migrated among organs, and the distribution of organs along the tips of B-cell lineage trees can provide important information about the nature of these cellular migrations. Visual inspection of the largest clonotypes in each mouse showed that in cohort-1x mice, clonotypes were typically restricted to either left or right lymph nodes, or shared among the spleen and bone marrow (**Fig. 4A** and **fig. S7A**). By contrast, large clonotypes in cohort-3x mice were more evenly dispersed among all six organs. These results are consistent with prior results showing a high degree of clonal overlap among left lymph nodes, right lymph nodes, and spleen / bone marrow in cohort-1x mice, with more widespread clones across all organs in cohort-3x mice (**Fig. 3** and **fig. S4**).

**Fig. 4.**
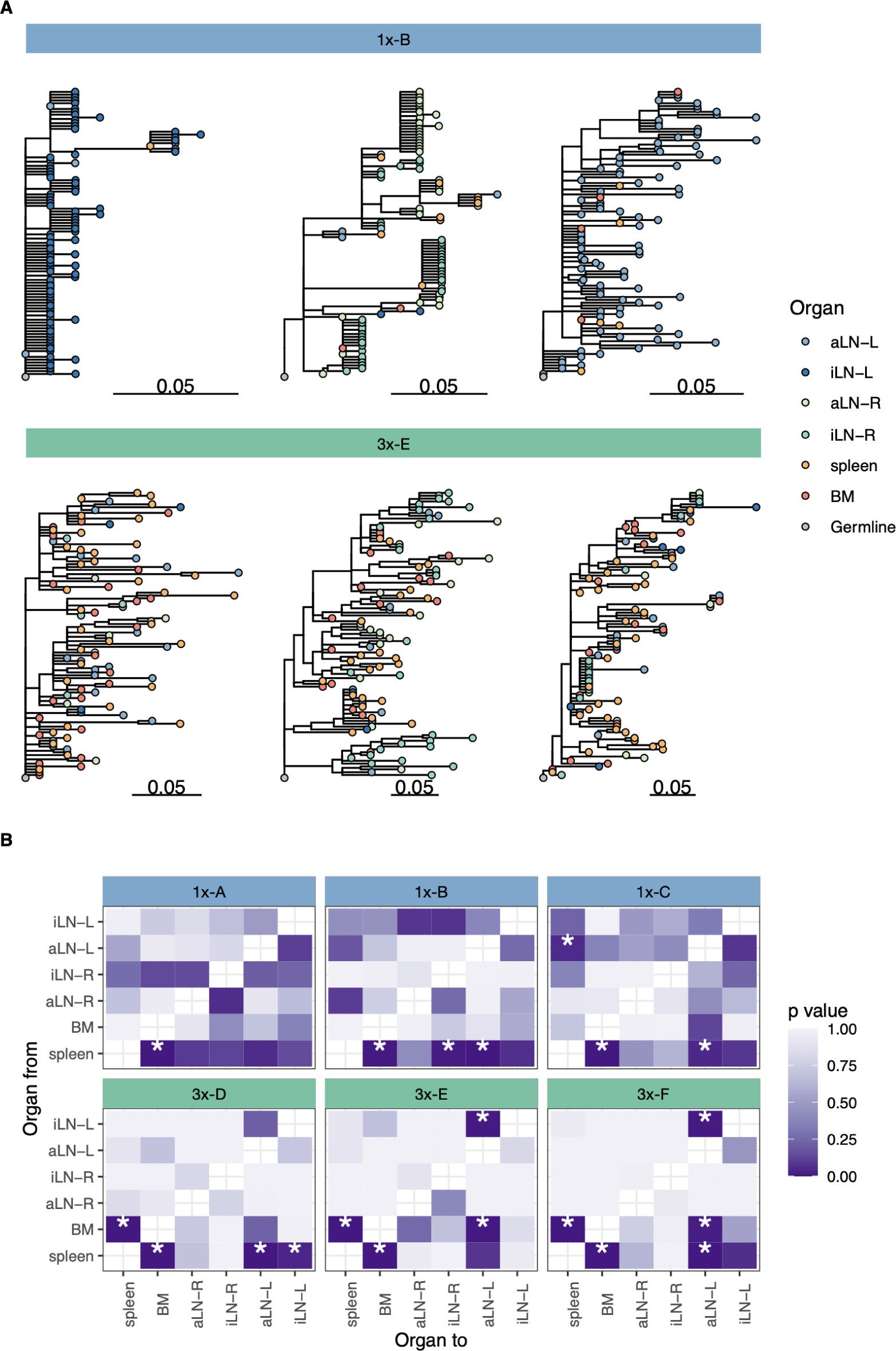
Phylogenetic analysis reveals physiological axes, diversification and sharing of clonotypes across multiple lymphoid organs. (**A**) Phylogenetic trees of the three most diverse clonotypes from a representative mouse from the single (1x-B) or multiple (3x-E) immunization cohorts. Color of nodes indicates the lymphoid organ origin of each tip with grey nodes depicting the clonotype’s unmutated germline sequence. Scale bar shows somatic hypermutations per codon site. See **Fig. S7A** for phylogenetic trees of the other cohort-1x and cohort-3x mice. (**B**) Heatmaps displaying p values derived from switch proportion (SP) tests, which quantify enrichment of transitions from one organ to another within lineage trees. BM: bone marrow, aLN-L, -R: left and right axillary lymph nodes, iLN-L, -R: left and right inguinal lymph nodes.

To further investigate the spread of clonotypes among lymphoid organs, we used the recently developed switch proportion (SP) test, which compares the proportion of each type of organ transition along observed trees to those obtained from trees with randomly assigned organ locations (*24*). A greater than expected proportion of transitions in a particular direction along lineage trees can be indicative of cellular migration patterns. We found that lineage trees in all cohort-1x mice had a significantly greater proportion of transitions from spleen to bone marrow than expected in random switching (**Fig. 4B**). By contrast, we found that lineage trees of all cohort-3x mice had a significantly higher proportion of transitions both from spleen to bone marrow, and from bone marrow to spleen. This indicates a close association between these organs in cohort-3x trees that is not biased in a particular direction. Repeating these analyses using only clonal sequences from spleen and bone marrow confirmed these patterns: cohort-1x mice had significantly more transitions from spleen to bone marrow, while cohort-3x mice did not (**Fig. S7B**). To detect associations between lymphoid organs within lineage trees, we repeated the SP test but permuted organ labels among trees within each mouse and quantified transitions in either direction between tissues. Consistent with prior results, we found a significantly greater proportion of switches between spleen and bone marrow in all mice (**Fig. S7C**). We also found a distinct left/right axis among lymph nodes, with all mice having a significantly greater proportion of transitions among left axillary and inguinal lymph nodes, as well as between right side lymph nodes (**Fig. S7C**). Overall, these phylogenetic analyses provide further evidence that B-cell clonotypes are more segmented along left, right and spleen / bone marrow axes upon a single immunization [at the left lateral flank], but are more widely dispersed following multiple immunizations.

### Physiological consolidation of antibody repertoires correlates with antigen-specificity

After uncovering that a strong humoral response in cohort-3x mice induced physiological consolidation of antibody repertoires, we next aimed to determine how this finding translates to antigen-specificity. Interrogating antigen-specificity on a repertoire-scale requires experimental screening platforms capable of linking antibody genotype and phenotype (antigen-binding) (*25–30*). Therefore, we utilized yeast surface display (*31*), fluorescence activated cell sorting (FACS) and deep sequencing to identify antigen-specific clones present in antibody repertoires (**Fig. 5A**). We generated three single-chain variable fragment (scFv) libraries from each of the cohort-3x mice by combinatorial (random) pairing of VH and variable light chain (VL) genes derived from bone marrow (**Fig. 5A, Fig. S8, A** and **B**). As there was a high degree of clonal overlap across all organs in cohort-3x mice, we decided that bone marrow would be a representative organ while also providing ample material for scFv library generation due to the presence of a high fraction of plasma cells expressing large amounts of IgG RNA (*32, 33*). scFv yeast display libraries were enriched for RSV-F antigen-binding by FACS over multiple rounds (**Fig. S8C**). By using a stringent FACS gating strategy, we ensured maximal purity at some expense of diversity, presumably by filtering out low-affinity clones. The final sorted population of yeast cells with specificity to antigen were subjected to deep sequencing of VH genes and the resulting CDRH3s were matched with the repertoire data of all organs within each corresponding mouse in order to assign RSV-F-specificity to antibody clones (now referred to as ‘binding clones’).

**Fig. 5.**
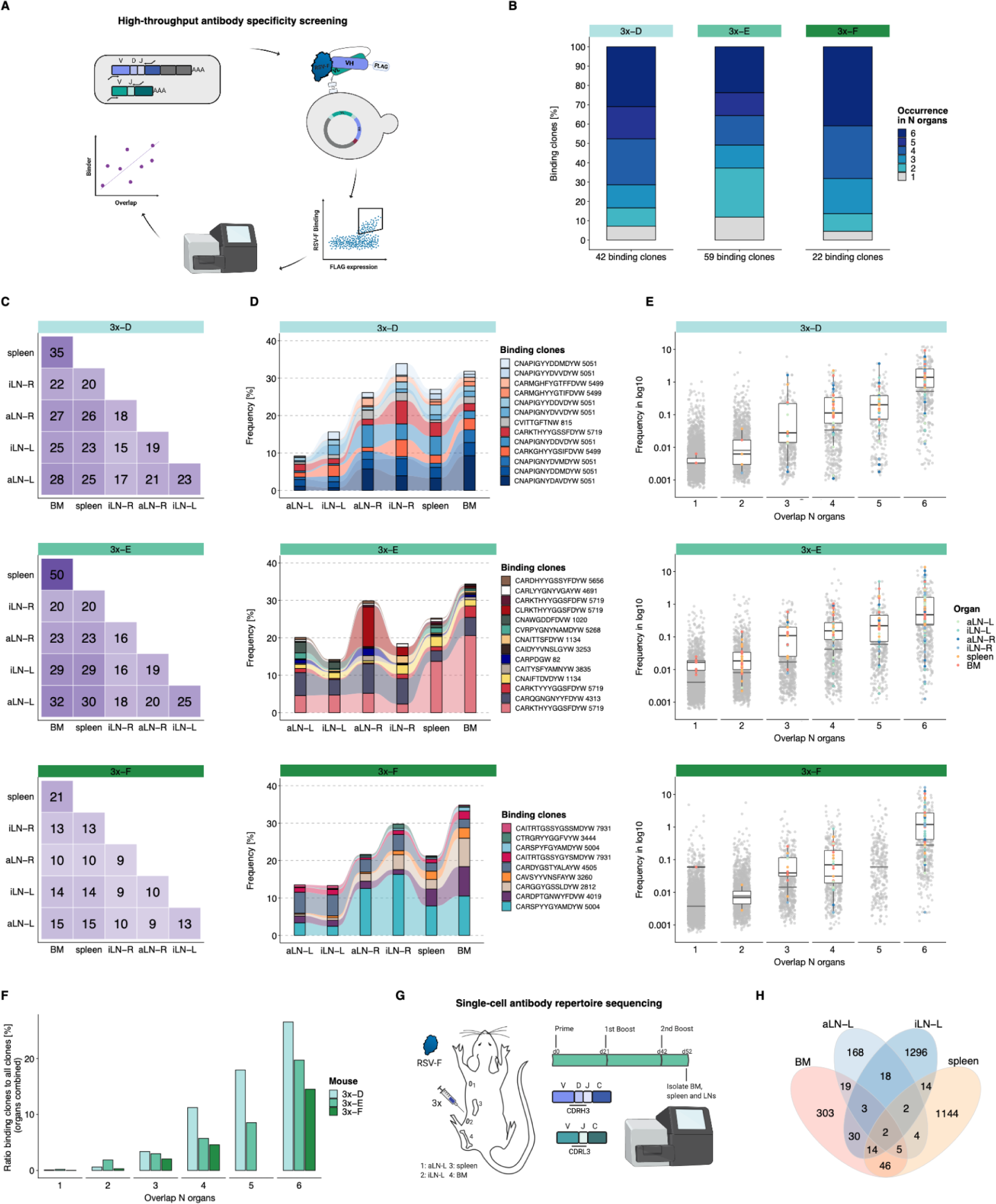
High-throughput antibody screening reveals that clonal overlap across lymphoid organs is correlated with antigen-specificity. (**A**) Workflow of high-throughput antibody screening by yeast display: antibody (scFv) libraries are generated by combinatorial pairing of VH and VL genes derived from bone marrow of cohort-3x mice and expressed on the surface of yeast cells, followed by isolation of antigen-binding cells by FACS and identification of clones by deep sequencing and repertoire analysis. (**B**) Bar plots represent proportions of binding clones occurring in *N* (1-6) organs within each corresponding mouse of cohort-3x (3x-D, 3x-E and 3x-F). (**C**) Heatmaps depicting numbers of shared binding clones between two organs within each mouse. (**D**) Tracking of binding clones shared across all six lymphoid organs and their relative frequencies for each mouse. Legend displays CDRH3 of each clone with the number indicating the corresponding clonotype group. (**E**) Distribution of clonal frequencies per degree of organ overlap (1-6) for all clones (grey) and binding clones (colored). Grey horizontal lines represent the median frequency of all clones, black lines including box plots represent the median frequency of binding clones within the corresponding overlap sections (see color legend for organ origin). (**F**) Bar plots displaying ratios of binding clones to all clones with regard to their degree of organ overlap (1-6) for each mouse. (**G**) Workflow for RSV-F immunizations into the left lateral flank of one Balb/c mouse, followed by single-cell antibody repertoire sequencing (VH + VL) of lymphoid organs (bone marrow (BM), spleen, left axillary and inguinal lymph nodes (aLN-L, iLN-L). (**H**) Venn diagram depicting numbers of shared unique IgG+ clones (based on identical V- and J- genes and CDRH3 + CDRL3) across lymphoid organs. BM: bone marrow, aLN-L, -R: left and right axillary lymph nodes, iLN-L, -R: left and right inguinal lymph nodes.

This workflow resulted in the detection of 120 unique binding clones from the three mice in cohort-3x, with the following distribution of binding clones (clonotypes) in each mouse: mouse 3x-D = 42 (13), 3x-E = 59 (23) and 3x-F = 22 (15). Importantly, 88-95% of these binding clones were shared between at least two organs out of which 24-41% were overlapping across all six lymphoid organs within the corresponding mouse (**Fig. 5B**). Since binding clones were experimentally derived from bone marrow RNA, the fact that the vast majority of clones (74-86%) were also observed in spleen confirmed the high degree of repertoire similarity in these two organs; additionally, a substantial fraction of binding clones was also observed in lymph nodes (51-68%) (**Fig. 5C)**.

Strikingly, the binding clones that were shared across all six organs occupied on average 24%, 24% and 22% of the corresponding organ repertoire in each mouse, respectively (**Fig. 5D**). When quantifying all clonal variants of clonotypes that were associated with the binding clones, left lymph nodes displayed comparable or even higher numbers of unique clonal variants than right lymph nodes, whereas the spleen and bone marrow showed consistently the highest numbers of clonal variants (**Fig. S9A**). Of note, we detected a shared (public) binding clonotype present in all six lymphoid organs of all three mice, thus hinting towards physiological consolidation and sequence convergence of antibody repertoires (**Fig. S9B**).

By also accounting for clonal expansion (clonal frequencies), we observed that the extent of organ sharing in binding clones positively correlated with higher clonal frequencies; i.e., clones shared across more organs were more clonally expanded (**Fig. 5E**). Finally, by calculating the ratio of binding clones to all clones, we could show a clear positive correlation between organ overlap and antigen-specificity, where 15-27% of shared clones across all organs could be confirmed as antigen-binding clones (**Fig. 5F**).

While our combinatorial scFv yeast display libraries could positively detect a substantial number of antigen-binding clones, it comes with the experimental limitation of random pairing of VH and VL chains. Therefore, in order to reconstruct the natural pairing of VH and VL chains, we performed single-cell antibody repertoire sequencing (10x Genomics, see methods) on IgG+ B cells and plasmablasts/plasma cells from lymphoid organs (bone marrow, spleen and left axillary and inguinal lymph nodes) of a mouse with a strong humoral immune response from receiving multiple booster immunizations (**Fig. 5G** and **fig. S10A**). Single-cell antibody sequencing recovered a range of 443-3416 IgG+ B cells per organ (**Fig. S10B**). Clonal repertoire analysis was performed using both VH and VL chains and clones were defined by identical germline V- and J-genes and 100% a.a. identity of CDRH3-CDRL3. This revealed once again substantial clonal overlap across lymphoid organs, including 26 clones shared across at least three organs and 157 clones shared in at least two organs (**Fig. 5H**). Antibody (scFv) expression and binding experiments confirmed that a substantial fraction (33%) of single-cell clones shared across organs were indeed antigen-specific (**Table S4** and **Fig. S10C**).

## Discussion

A central pillar of antibody repertoires is their inherently high diversity (estimated to be >10^13^ in mice) (*17*). Therefore, each lymphoid organ may be populated with an exclusive repertoire of naive B cells, which following a single immunization will induce independent B-cell activation, diversification and germinal center reaction events. To investigate how these dynamic processes shape antibody repertoires across lymphoid organs during an immune response, we performed deep sequencing of antibody repertoires using two cohorts of mice that were immunized either once or three times.

Analysis of antibody repertoires in the single immunization cohort revealed extensive clonal expansion in lymph nodes from both left and right sides, suggesting antigen-induced selection and expansion of an IgG-class-switched subset of B cells. These may be comprised of early germinal center B cells and short-lived plasmablasts, the latter being generated within the first days after an immunization or infection and are considered as transient precursors of plasma cells (*34–36*). A recent study reported similar findings of highly expanded germinal center B cells in draining lymph nodes seven days after influenza infection in mice (*10*). Overlap analysis of the most expanded clones revealed distinct physiological axes: inguinal and axillary lymph nodes from the left side shared the majority of their expanded clones, whereas the same was true for right-side lymph nodes, with minimal overlap across left and right lymph nodes. This may reflect migration of antigen-activated B cells and/or antigen transport across the circulatory network, and lymphatic connections between inguinal and axillary lymph nodes have been well described (*37, 38*). A majority of clones that were expanded in lymph nodes were also found in spleen and bone marrow, however, exhibiting lower frequencies. This may indicate early homing of B cells to the spleen and bone marrow where they reside as terminally-differentiated plasma cells (*5, 14*). This is supported by phylogenetic tree analysis showing significantly more transitions from spleen to bone marrow in single-immunized mice. Performing analysis on all clones, however, revealed limited clonal diversification and low overlap across all lymphoid organs, demonstrating that a primary and weak humoral response is characterized by clonal expansion and segregation of a few clones to particular organs while exhibiting mostly lymphoid organ-exclusive repertoires.

Strikingly, a strong humoral response resulted in a substantial number of shared clones in all mice, exhibiting an up to 70-fold increase in clonal overlap across all six lymphoid organs. Global repertoire similarity measurements confirmed high degrees of pairwise clonal overlap between organs, with shared clones exhibiting similar frequencies. This is in line with the recent observation of a close association of distinct lymphoid tissues that form B-cell clonal networks in humans (*7*). Two mechanisms may be contributing to this high clonal overlap and repertoire similarity in lymphoid organs: (i) subcutaneous injection of soluble antigen results in drainage from interstitial space into lymphatic vessels and is transported directly to lymph nodes (*32, 39*), with eventual transport into systemic circulation, including the spleen (*40*); (ii) migration of antigen-presenting cells (e.g., dendritic cells) (*41*) and antigen-specific memory B cells via lymphatic and systemic circulation (*42*). Network-based analysis revealed extensive clonal diversification, which is consistent with the refueling of ongoing germinal center reactions (which usually subside after 30 days) (*11*). This supports antigen transport to all secondary lymphoid organs and B cells, re-entering or still residing in germinal centers, undergoing multiple rounds of selection and affinity maturation, which was reflected by high levels of somatic hypermutation across all organs. Moreover, phylogenetic analysis was performed to investigate if physiological consolidation across lymphoid organs was a consequence of B-cell migration and indeed revealed significant clonal lineage transitions between spleen and bone marrow as well as from spleen to left lymph nodes (side of immunization).

By performing high-throughput screening using yeast surface display, we confirmed that a major fraction of shared clones across organs was antigen-specific. Also, we observed that spleen and bone marrow exhibited the highest repertoire similarity, by sharing the majority of their detected binding clones. Moreover, we uncovered that shared binding clones displayed relatively high clonal frequencies, which is in line with reports showing that antigen-specific serum titer responses are mainly a result of a few expanded B-cell clones, reflecting oligoclonal B-cell responses upon vaccination or infection (*32, 43, 44*).

Here, we have systematically uncovered that strong humoral responses induce a high degree of physiological consolidation of antibody repertoires and antigen-binding clones, observed across multiple distinct lymphoid organs, which is also associated with an overall reduced clonal diversity. Although these findings are dependent on immunization parameters [i.e., antigen delivery (viral-based, protein, mRNA) and adjuvant types], they provide important implications for vaccinology. For example, multiple immunization schemes are often employed clinically (e.g., mRNA vaccinations for Covid-19 and hepatitis B subunit vaccines) (*45, 46*) to induce strong humoral responses (based on antigen-binding serum titers), however, this may come at the expense of reducing antibody repertoire diversity across lymphoid organs. Future vaccine strategies may benefit by attempting to balance the strength of the induced humoral response with the physiological landscape of clonal repertoire diversity, as more diverse antibody repertoires in lymphoid organs may facilitate greater breadth of antigen coverage and perhaps enhance resistance to viral escape mutations (e.g., SARS-CoV-2 and influenza).

## Materials and Methods

### Study design

Here, we aimed to interrogate the physiological landscape of antibody repertoires upon immunization. We performed an in-depth systems analysis that included murine models of immunization, antibody repertoire sequencing, bioinformatic repertoire analysis and high-throughput antibody specificity screening. The study was designed around two cohorts of mice that either were immunized once (cohort-1x, n=3) or three times (cohort-3x, n=3), resembling a weak and strong immune response, respectively, which was confirmed by serum titer ELISA. We performed deep sequencing on IgG antibody repertoires of multiple lymphoid organs: bone marrow, spleen and left and right axillary and inguinal lymph nodes. Bioinformatic analysis focused on assessment of clonal diversity, clonal expansion, repertoire similarity (based on clonal overlap and germline V-gene usage), and clonal diversification (phylogenetic and sequence-similarity network analysis) across all organs within each mouse. For assessment of antigen-specificity across organ repertoires, we generated bone marrow-derived combinatorial scFv yeast libraries of cohort-3x mice. Additionally, we performed single-cell antibody repertoire sequencing (10x Genomics) of lymphoid organs (bone marrow, spleen and left axillary and inguinal lymph nodes) from one mouse that was immunized three times.

### Mouse immunizations and lymphoid organ harvest

All mouse experiments were performed under the guidelines and protocols approved by the Basel-Stadt cantonal veterinary office (Basel-Stadt Kantonales Veterinäramt, Tierversuchsbewilligung #2582). Six female Balb/c mice (Janvier Laboratories France, 10 weeks old) were housed under specific pathogen-free conditions and maintained on a standard chow diet. Mice were separated into two cohorts of three mice and immunized subcutaneously into the left flank with 10 µg respiratory syncytial virus (pre-)fusion glycoprotein (RSV-F, expressed and purified as previously described (*47*)) in combination with 50 µg poly(I:C) adjuvant (HMW, InvivoGen tlrl-plc) per mouse. RSV-F protein was resuspended in 1X DPBS (Gibco 14190144) together with poly(I:C) adjuvant to a final injection volume of 100 µl per mouse shortly before each immunization. One cohort received a single immunization (cohort-1x), whereas the other cohort received three immunizations with 21 days intervals between booster immunizations (cohort-3x). Mice from both cohorts were euthanized by CO_2_ asphyxiation and cervical dislocation ten days after the single or third immunization, respectively. From each mouse, left and right axillary lymph nodes, left and right inguinal lymph nodes, spleen, and both femurs and tibiae were harvested. In order to avoid contamination, pipettes and scissors were cleaned and sterilized after each organ isolation step. Blood samples were collected by cardiac puncture, followed by separation of serum after 30 minutes clotting at room temperature and two subsequent centrifugation steps at 6000g for 15 minutes. Lymph nodes and spleens were directly placed in 1-2 ml RNAlater (Sigma R0901) and stored at 4°C for up to six days and transferred to -20°C until further processing. Isolated femurs and tibiae were stored in a buffer (PBS, containing 0.5 % BSA (Miltenyi Biotec 130-091-376) and 2 mM EDTA (Thermo Fisher Scientific AM9260G) on ice. For bone marrow isolation, both ends of femurs and tibiae were cut with surgical scissors and bone marrow was extracted by flushing 3-5 ml sterile and ice-cold buffer through the bones using a 26G needle (BD Microlance 3 300300). The isolated bone marrow was then filtered through a 40 µM nylon cell strainer (FALCON 352340) and cell-suspension was centrifuged at 300 g for 10 minutes at 4°C. The supernatant was removed and 1 ml of Trizol (Life Technologies 15596) was used to resuspend samples before storing at -80°C until further processing.

For the single-cell sequencing experiment, the same procedure as for cohort-3x mice was done for one female Balb/c mouse (Janvier Laboratories France, 9 weeks old). At the end of the experiment, left-side axillary and inguinal lymph nodes, as well as spleen and both hindlegs were isolated and stored in freshly prepared and ice-cold buffer (PBS + 2% Fetal Bovine Serum (FBS) (Thermo 16250078) + 2mM EDTA, sterile filtered). Bone marrow was isolated as described above and kept on ice until further processing for FACS.

### RNA isolation from lymphoid organs

Mouse spleens and lymph nodes were removed from RNAlater solution and transferred to gentleMACS M tubes (Miltenyi Biotec 130-093-236) and 2 ml microtubes (OMNI 19-629-3, 1.4 and 2.8 mm Ceramic mix) containing 1.5 ml and 0.8 ml ice-cold Trizol, respectively. Spleens and lymph nodes were dissociated using gentleMACS Octo Dissociator (Miltenyi Biotec 130-095-937, RNA_01_01 pre-programmed setting was performed twice) and Bead Ruptor 24 Elite (OMNI 19-040E, 1x 10 sec, 4 m/s, 1 cycle), respectively. Cell-suspensions were removed and stored at -80°C (spleen) or subsequently processed for RNA extraction (lymph nodes). All samples were thawed on ice prior to RNA extraction using the PureLink RNA Mini Kit (Life Technologies 12183018A) following the manufacturer’s guidelines. Briefly, 0.7 ml of lymph node cell suspension and 1 ml of spleen and bone marrow samples were loaded with chloroform onto 5PRIME Phase Lock Gel Heavy tubes (Quantabio, 2302830) for better separation of phases. All other steps were performed according to the manual’s instructions (protocol: Using Trizol Reagent with PureLink RNA Mini Kit). Samples were eluted in 30 / 80 µl water (lymph nodes / spleen, bone marrow) and stored at -80°C until further processing. After RNA isolation, overall RNA quality was determined by RNA ScreenTape Analysis (Agilent, 5067-5576), with all samples showing RIN values between 8 and 10.

### Antibody repertoire library preparation and deep sequencing

Antibody variable heavy chain (VH) libraries for deep sequencing were constructed using a previously established protocol of molecular amplification fingerprinting (MAF), that incorporates unique molecular identifiers for error and bias correction (*19*). Briefly, first-strand cDNA synthesis was performed using Maxima reverse transcriptase (Life Technologies EP0742) following the manufacturer’s instructions, using 5 μg total RNA and a gene-specific primer corresponding to constant heavy region 1 of IgG subtypes (IgG1, IgG2a, IgG2b, IgG2c, and IgG3) with an overhang region containing a reverse unique molecular identifier (RID). After cDNA synthesis, samples were subjected to a left-sided 0.8X SPRIselect bead cleanup (Beckman Coulter B23318). Quantification of target-specific cDNA by a digital droplet (dd)PCR allowed exact input of first-strand cDNA copies of each sample into the next PCR reaction. Multiplex-PCR was performed using a forward primer set annealing to framework 1 regions of VH including an overhang region of forward unique molecular identifier (FID) and a partial Illumina adapter as well as a reverse primer containing a partial Illumina sequencing adapter. The reaction mixtures contained 135.000 first-strand cDNA copies, 15.000 synthetic spike-ins (for quality control), primer mix, and 1X KAPA HIFI HotStart Uracil+ ReadyMix (KAPA Biosystems KK2802) with the Uracil+ version enabling efficient high-fidelity amplification of multiplex primers containing deoxyinosine. PCR reactions were cleaned using left-sided 0.8X SPRIselect bead clean-up as before and product was quantified again using ddPCR assay. Finally, an Illumina adaptor-extension singleplex PCR step was performed using 820.000 copies of the previous PCR product with 1X KAPA HIFI HotStart ReadyMix (KAPA Biosystems KK2601), followed by double-sided (0.5X–0.8X) SPRIselect bead cleanup and sample elution in tris-EDTA buffer (Fluka 93302-100ML). Overall library quality and concentration was determined on the Fragment Analyzer (Agilent DNF-473 NHS Fragment kit). Libraries were then pooled in Tris-HCl + 0.1% Tween-20 (Teknova T7724) and sequenced on an Illumina MiSeq using the reagent v3 kit (2x300 bp) with 10% PhiX DNA for quality purposes. All 36 samples were processed and sequenced in three batches of 12 samples by preparing sequencing libraries of one mouse per cohort per batch with mixed sample order.

### Antibody repertoire analysis

Raw FASTQ files were processed by a custom CLC Genomics Workbench 10 script. First, low quality nucleotides were removed using the quality trimming option with a quality limit of 0.05, followed by merging of forward and reverse read pairs and removal of sequences not aligning to mouse IGH constant sequences. Only amplicons between 350 and 600 base pairs were kept for further processing. Pre-processed sequences were then used as input for the previously established MAF bioinformatic pipeline 12 (*19*) to perform alignment of sequences as well as error and bias correction. For all downstream analysis, an output TSV file (‘CDR3_Tot_Table’) for each sample was used that listed each unique CDRH3 of the consensus-built data per row, including majority V-gene, majority J-gene information, corrected MAF frequency information, and somatic hypermutation statistics. Within the study, we defined sequences with identical CDRH3 a.a. sequence and matching V-, and J-genes to be a clone. For investigation of clonally related B-cell variants (here referred to as clonotypes), we performed hierarchical clustering (single linkage) of antibody sequences with identical V-, and J-gene and matching CDRH3 a.a. lengths and > 90% a.a. CDRH3 identity (*48*).

For phylogenetic tree analysis, we used the output TSV file ‘Polished_Annotated_Table’ containing all corresponding consensus built and error-corrected read sequences belonging to a clone (here referred to as clonal sequences) in order to obtain intraclonal variants for building lineage trees.

All downstream analysis was carried out using customized R scripts in R version 4.0.3 (*49*).

### Comparative analysis of germline V-gene usage

Counts for unique germline V-genes were summarized for all unique clones within each sample (without including clonal frequency information) and Pearson’s correlation coefficients were calculated between two repertoires across all samples.

### Quantification of clonal expansion

To assess the distribution of clonal frequencies in the repertoires, we quantified clonal expansion profiles (Hill-based Evenness profiles) as previously described (*20*). Hill-based Evenness profiles indicate the state and uniformity of clonal expansion by using a continuum of diversity indices (the parameter alpha determines the impact of high frequency clones: with increasing alpha values, higher frequency clones are weighted more).

### Assessment of repertoire similarities

Analysis of pairwise repertoire similarities were performed by quantification of pairwise Jaccard and cosine similarity using R package immunarch 0.6.5 (*50*). Briefly, we calculated the pairwise Jaccard index, by quantifying the size of intersection between two repertoires A and B and dividing it by the length of the union of the same repertoires, to assess repertoire similarities based on clone overlap across all samples: 

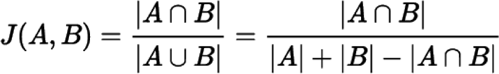

Cosine similarity measurements indicate the degree of antibody repertoire similarity between two repertoires A and B by including both clonal overlap and frequency information (by giving weight to clonal frequencies), ranging from 0 to 1, thus higher cosine values correspond to higher similarities between two repertoires (*7*): 

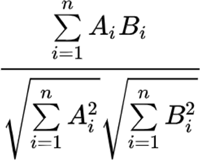

Additionally, overlapping unique clones were represented graphically via venn diagrams per mouse and cohort using R package venn 1.9 (*51*).

### Network-based analysis

Sequence-similarity networks of clonotypes were constructed as described in the figure legends. Briefly, each node represents a unique clone that is connected by edges to clones with a Hamming distance of 1 between CDRH3 a.a. (edit distances were calculated using the R package stringdist v.0.9.6.3 (*52*)). The relative size of each node was proportional to its clonal frequency (in log and rescaled for each mouse). Networks were visualized with the Kamada-Kawai layout; all network and connectivity plots were generated using the R package igraph v.1.2.6 (*53*).

### Phylogenetic tree analysis

Phylogenetic analysis of BCR sequences is best performed using full-length BCR sequences. Full length V region sequences were obtained from MAF output TSV file ‘Polished_Annotated_Table’. Only sequences that were productively rearranged, associated with sequences in the final CDRH3 dataset, and did not correspond to spike-in control sequences were retained for final analysis. Within each sample, identical sequences were collapsed and sequences with less than 3 associated reads were discarded using *presto* v0.6.2 (*54*). Sequences were aligned to the IMGT GENE-DB v3.1.24 (*55*) mouse immunoglobulin database using IgBlast v1.13.0 (*56*). Sequences from all tissue samples within each mouse were clustered into clones by partitioning based on common IGHV gene annotations, IGHJ gene annotations, and junction lengths. Within these groups, sequences differing from one another by a length normalized amino acid Hamming distance of within the junction region were defined as clonotypes by single-linkage clustering using *scoper* v1.1.0 (*57*). This Hamming distance threshold and use of amino acid Hamming distance was used for consistency of clonotype definition in other analyses within this manuscript.

Phylogenetic analysis was performed using *dowser* v0.1.0 (*24*). Within each mouse, sequences from the same tissue differing only by ambiguous nucleotides were collapsed. For computational efficiency, large clones were randomly down-sampled to a maximum size of 100 sequences. Clones containing sequences from only one organ, or fewer than 20 sequences total, were removed. Lineage tree topologies and branch lengths were estimated using *IgPhyML* v1.1.3 (*58*) and visualized using *ggtree* v2.4.2 (*59*). The “switch proportion” (SP) permutation test (*24*) was used to assess whether lineage trees showed signs of migration among organs in a particular direction. Briefly, given the set of organ labels at the tips of each phylogenetic tree, a maximum parsimony algorithm implemented in *IgPhyML* was used to reconstruct the set of internal node organ labels resulting in the fewest number of organ changes along the tree. For all trees within each mouse, the number and direction of organ changes along all trees was recorded and normalized by the total number of changes to give the switch proportion statistic (SP). Organ states were then randomized within each tree, and the resulting SP statistic was calculated for these permuted trees. The difference between observed and permuted SP statistics (δ) was recorded, and this process was repeated for 1000 replicates. The P value for enrichment of changes between tissues (i.e., δ > 0) is the proportion of replicates in which δ ≤ 0. If P < 0.05 and δ > 0 for a given pair of tissues, this indicates a significantly more biased ancestor/descendant relationship from one tissue to the other than expected by chance in the lineages surveyed. Tree topologies were not re-estimated throughout permutation replicates; however, to control the false positive rate of the SP test, all trees were randomly down-sampled to a maximum tip-to-state change ratio of 20 for each repetition. Clusters of internal nodes separated by zero length branches (polytomies), were re-ordered using nearest-neighbor interchange (NNI) moves to minimize the number of changes along the tree and to appropriately represent possible directions of migration. For analysis in **Fig. S7C**, the SP test was repeated as above but node organ labels were permuted among trees within each mouse. Further, tissue changes were quantified in either direction among tissues. This modified SP test more directly quantifies association among organs, rather than biased ancestor-descendant relationships.

### Antigen ELISA

Standard antigen enzyme-linked immunosorbent assays (ELISAs) were performed to measure mice sera for RSV-F specific titers. High binding 96-well plates (Costar CLS3590) were coated overnight with 2 µg/ml of RSV-F in PBS at 4°C. Plates were washed three times in PBS containing 0.05% (v/v) Tween-20 (AppliChem A1389) (PBST) and blocked for two hours at room temperature with PBS containing 2% (m/v) non-fat dried milk powder (AppliChem A0830) and 0.05% (v/v) Tween-20 (PBSM). After blocking, plates were washed three times with PBST. Sera were prediluted depending on cohort (1:150 for cohort-1x and 1:450 for cohort-3x) and then serially diluted in 1:3 steps in PBSM across the plate. For normalization of multiple plates, motavizumab, a humanized monoclonal RSV-F binding antibody, was used as a standard curve by serial 1:5 dilutions (starting concentration: 1 µg/ml). Plates were incubated for one hour at room temperature and washed three times with PBST. HRP-conjugated rat monoclonal [187.1] anti-mouse kappa light chain antibody (abcam ab99617, 1:1500 dilution in PBSM) and HRP-conjugated goat polyclonal anti-human IgG F(ab’)_2_ fragment antibodies (Jackson ImmunoResearch 109-036-008, 1:5000 dilution in PBSM) were used as secondary detection antibodies for sera and Motavizumab, respectively. Plates were incubated at room temperature for one hour again, followed by three washing steps with PBST. ELISA detection was performed using the 1-Step Ultra TMB-ELISA Substrate Solution (Thermo 34028) and reaction was terminated adding 1 M H_2_SO_4_. Absorption at 450 nm was measured with the Infinite M200 PRO NanoQuant (Tecan) and data were analyzed using Prism V7 (Graphpad). Serum endpoint titer was determined using the last diluted specimen that gave a positive signal compared to the negative control (no serum).

### Cloning and expression of yeast display antibody libraries

For each mouse of cohort-3x, two to three independent combinatorial scFv yeast display libraries were constructed using corresponding bone marrow RNA samples for detection of RSV-F binding antibody sequences (based on CDRH3). It should be noted that constructing a randomly paired VH VL library may not cover the whole repertoire space due to improper VH VL pairing.

Briefly, reverse transcription of bone marrow RNA samples (same as used for antibody repertoire sequencing) was performed using Maxima reverse transcriptase (Life Technologies EP0742) following the manufacturer’s instructions. Two 40 µl reactions per sample were prepared using 1 μg total RNA and IgG gene-specific primers (p1/p2, see **Table S5**) for heavy chains or oligo(dT)_18_ primer (Thermo, SO131) for light chains. After cDNA synthesis, split-pool PCR was performed for VH and VL samples by setting up eight parallel 25 µl reactions for each sample, containing 4 µl of first-strand cDNA, the VH or VL primer mix (reported in **Table S5**, final concentration of 0.5 µM each) and Kapa HiFi HotStart Uracil+ Ready Mix (Kapa Biosystems KK2802) or Kapa HiFi HotStart Ready Mix (Kapa Biosystems KK2601) for VH and VL reactions, respectively. The following cycling conditions were used for VH amplification: initial denaturation 3 min at 95°C, 30 cycles with denaturation at 98°C (20 s), annealing at 60°C (30 s), elongation at 72°C (45 s), and final elongation at 72°C (1 min); for VL amplification: initial denaturation 3 min at 95°C, 30 cycles with denaturation at 98°C (20 s), annealing at 54°C (15 s), elongation at 72°C (30 s), and final elongation at 72°C (1 min). After split-pool PCR, the corresponding eight reactions were pooled and subjected to a first PCR clean-up with QIAquick PCR Purification Kit (Qiagen 28016) to concentrate samples, followed by a final clean-up step by agarose-gel extraction (2% (w/v) gel) with Zymoclean Gel DNA Recovery kit (Zymo Research D4001). Yeast scFv display libraries were generated using the amplified VH and VL DNA libraries from above and the pYD1 yeast surface display vector (Addgene plasmid #73447; https://www.addgene.org/73447/; RRID:Addgene_73447) (*60*), which was modified to include a BamHI restriction site between HA- and FLAG-epitope tags (from now on referred to as ‘pYD1-BamHI’) [containing galactose (GAL) induced GAL1 promoter and a BamHI restriction site in order to insert both VH VL libraries separated by a (Gly_4_Ser)_3_ linker as scFv format] (**Fig. S8B**). All in-frame scFv sequences resulted in a C-terminal FLAG-tag to identify scFv-expressing yeast cells. The vector was linearized using BamHI-HF (NEB R0136S) in Cutsmart buffer (NEB B7204S) by incubation for 45 minutes at 37°C and immediate purification using Zymo DNA Clean & Concentrator kit (Zymo D4005). Shortly before transformation, inserts and cut vector were dialysed for 30 minutes using 0.025 µm MF-Millipore membrane filters (Merck VSWP01300) and purity of final products was confirmed by agarose gels and concentration measured using NanoDrop 2000c (Thermo Scientific ND-2000). Next, 8-10 separate reactions per library were prepared by combining 1µg of linearized vector with 1µg of each VH and VL inserts for co-transformation into 330 µl competent EBY100 yeast cells. Electroporation of EBY100 yeast cells was performed with BioRad MicroPulser Electroporator (BioRad 1652100) using pre-chilled Gene Pulser cuvettes (2mm gap, BioRad 1652086) as described previously (*61, 62*). Transformation efficiency was determined by plating tenfold serially diluted yeast cells on SD-CAA + 2% glucose plates following incubation at 30°C for 48 h. This workflow resulted in ∼1-3x10^6^ transformants per library. Transformation efficiency could be optimized by using longer homology overhangs in the primers used for VH and VL amplification. As positive control, monoclonal yeast cells expressing the RSV-F binding monoclonal antibody Palivizumab as scFv format were generated (see method section ‘Generation of monoclonal scFv yeast cells’).

### Screening yeast display antibody libraries for antigen binding by FACS

Yeast cells were cultured in yeast nitrogen base-casamino acids (YNB-CAA) (BD Biosciences 223120) + 2% D-(+)-Glucose (Sigma G5767-500G) growth medium including phosphate buffer (5.4 g/l Na_2_HPO_4_, 8.6 g/l NaH_2_PO_4_·H_2_O) and 1X Penicillin-Streptomycin (P/S) (Life Technologies 15140122) to avoid bacterial contamination. A day before sorting, a fraction of yeast cells, > 10-fold larger than the (initial or sorted) library sizes, were pelleted at 3.000g for 3 minutes and induced by resuspension in YNB-CAA + 2% D-(+)-Galactose (Sigma G0625-500G) + phosphate buffer + P/S induction medium at a final OD_600_ of 0.5. Cells were grown at 30 °C overnight, with shaking at 250 rpm. On the day of sorting, a total of 10 - 50-fold larger than the (initial or sorted) library sizes were pelleted at 7.000 g for 2 minutes at 4°C, washed twice in buffer (PBS + 0.5% BSA) and stained with the anti-FLAG-phycoerythrin (PE) antibody (anti-DYKDDDDK Tag, BioLegend 637310) for confirmation of scFv expression and RSV-F protein conjugated with Alexa Fluor 647 (AF647) to select antigen binding cells. For the antibody labeling steps, yeast cells were resuspended in ice-cold buffer containing 0.02 μg/μL anti-FLAG-PE and 0.01 μg/ μL RSV-F-AF647 with a final cell density of ∼10^5^ cells/μL. Cells were incubated for 30 minutes at 4°C with shaking at 450 rpm. Finally, cells were washed twice and filtered before flow sorting. The following machines were used: Sony SH800S (Sony Biotechnology), Sony MA900 (Sony Biotechnology), or BD FACS Aria III (BD Biosciences). Flow cytometry data was analyzed using FlowJo V10.4.2 (FlowJo, LLC).

For setting the sort gates, monoclonal yeast cells expressing Palivizumab as scFv were used as positive control and corresponding libraries stained with anti-FLAG-PE only were used as negative controls. Cells were sorted for double-positive cells (AF647+/PE+), and recovered yeast cells were cultured for expansion in glucose growth medium. The expansion, induction, staining, and recovery were performed three additional times with the same antigen and working concentration. Gates were set less stringent due to the low fraction (∼0.1-0.3%) of double-positive cells for the first sort, but stringency was increased with further sort rounds. After the final sort, the enriched cells were screened for purity of double-positive yeast populations using BD LSR Fortessa (BD Biosciences) and enriched yeast plasmid libraries were extracted using Zymoprep Yeast Plasmid Miniprep II kit according to the manufacturer’s instructions (Zymo Research D2004).

### Deep sequencing of antigen-binding yeast display libraries

The purified plasmid DNA was used as a template for PCR using a primer pair (LC62 and LC63, see **Table S5**) targeting the VH of the recombined scFv insert for detection of RSV-F binding antibody sequences (based on CDRH3). Briefly, per library two 25 µl reactions were prepared using 2 µl of plasmid template each together with p50 and p51 primers (final concentration of 0.8 µM) and Kapa HiFi HotStart Ready Mix. The following cycling conditions were used: initial denaturation 3 min at 95°C, 23 cycles with denaturation at 98°C (20 s), annealing at 65°C (30 s), elongation at 72°C (45 s), and final elongation at 72°C (1 min). Samples were subjected to a left-sided 0.8X SPRIselect bead cleanup and an Illumina adaptor-extension PCR step was performed using 150 ng of purified PCR product into 50 µl singleplex PCR with 1X KAPA HIFI HotStart ReadyMix and Illumina Nextera primers for dual indexing, followed by double-sided (0.5X–0.8X) SPRIselect bead cleanup and sample elution in tris-EDTA buffer. Overall library quality and concentration was determined on the Fragment Analyzer (Agilent DNF-473 NHS Fragment kit). All libraries were then pooled and sequenced on an Illumina MiSeq using the reagent v3 kit (2x300 bp) with 20% PhiX DNA for quality purposes.

Raw sequencing files obtained from Illumina MiSeq were aligned using the “analyze amplicon” command of the MiXCR software package (*63*). For downstream analysis, sequences with a clone count of at least 2 were kept and all unique CDRH3s obtained from libraries of the corresponding mouse were pooled. This CDRH3 pool was then matched with the corresponding mouse repertoire data of all six lymphoid organs in order to assign RSV-F-specificity to antibody clones (referred to as ‘binding clones’).

### FACS isolation of B-cell subsets

We performed fluorescence activated cell sorting (FACS) to isolate IgG+ CD19+ B cells from lymph nodes and spleen as well as plasma cells and plasmablasts (CD19+, TACI+, CD138+) from spleen and bone marrow (as previously described (*64*)) in order to generate antibody repertoire libraries for single-cell sequencing.

First, single-cell suspensions were prepared from spleen, left axillary and left inguinal lymph nodes by transferring each organ and ice-cold buffer (DPBS, 2 % FBS and 2 mM EDTA) onto a 40 µM nylon cell strainer and softly mashing the organ through the cell strainer into a petri dish using the plunger tip of a 1 ml syringe (BD Plastipak 303172). Cell suspensions were filtered several times and filters were rinsed with fresh buffer to obtain a homogeneous single-cell suspension. All steps were performed on ice if not stated otherwise and all centrifugation steps were performed at 300 g for 5 minutes at 4°C.

For spleen and bone marrow suspensions, red blood cell lysis was performed after centrifugation of cell suspensions, followed by resuspension and incubation in 3 / 5 ml RBC lysis buffer (eBioscience 00-4333-57) for 5 min at room temperature. Lysis was stopped by adding 10 ml of ice-cold buffer and filtering of cell-suspension through a 40 µM nylon cell strainer. Cells were counted and 5E7 cells per spleen and bone marrow, and 5 - 8E6 cells per lymph nodes were used centrifuged for each staining. To avoid nonspecific binding, cell pellets were resuspended in x μL of unlabeled CD16/32 mAb Purified Rat Anti-Mouse CD16/CD32 (Mouse BD Fc Block™), BD Pharmingen 553141), 1:50 in PBS + 2% FCS) and blocked for 15 min on ice. Next, cells were washed in ice-cold buffer and centrifuged before staining. The pellet was then resuspended to a final concentration of 5E6 cells/ml in buffer containing the respective fluorochrome-coupled antibodies and incubated for 30 min on ice in the dark. The following antibodies were used: Brilliant Violet 421 anti-mouse IgD Antibody (BioLegend 405725, 1:100 dilution), Brilliant Violet 421 anti-mouse IgM Antibody (BioLegend 406517, 1:100 dilution), APC-Cyanine7 anti-mouse CD4 Antibody (BioLegend 100413, 1:200 dilution), APC-Cyanine7 anti-mouse CD8a Antibody (BioLegend 100713, 1:200 dilution), APC-Cyanine7 anti-mouse NK-1.1 Antibody (BioLegend 108723, 1:200 dilution), APC-Cyanine7 anti-mouse TER-119 (BioLegend 116223, 1:200 dilution), PE-Cyanine7 anti-mouse CD19 (BioLegend 115519, 1:200 dilution), FITC anti-mouse IgG1 (BioLegend 406605, 1:100 dilution), FITC anti-mouse IgG2a/2b (BD Pharmingen 553399, 1:100 dilution), FITC anti-mouse IgG3 (BD Pharmingen 553403, 1:100 dilution), PerCP-Cyanine5.5 anti-mouse CD138 (BioLegend 142509, 1:200 dilution), PE anti-mouse CD267 (TACI) (BioLegend 133403, 1:200 dilution). Cells were incubated with the antibody mix including LIVE/DEAD Fixable Near-IR Dead Cell Stain (1:10.000, Life Technologies L34975) for 30 minutes at 4°C. Cells were washed twice and resuspended before cell sorting into ice-cold buffer using a FACSAria with FACSDiva software.

### Single-cell sequencing and analysis of antibody repertoires

Single-cell sequencing libraries were prepared from sorted cells according to the provided 10x Genomics’ protocol (‘Direct target enrichment - Chromium Single Cell V(D)J Reagent Kits’ (CG000166 REV A)). Briefly, sorted single cells were co-encapsulated with gel beads (10x Genomics, 1000006) using 4 lanes of one Chromium single cell chip by loading 1.474 - 13.000 cells per reaction. V(D)J library preparation was performed using the Chromium Single Cell 5’ Library Kit (10x Genomics 1000006) and the Chromium Single Cell V(D)J Enrichment Kit, Mouse B Cell (10x Genomics 1000072) according to the manufacturer’s manual. Final libraries were pooled and sequenced on the Illumina NextSeq 500 platform (mid output, 151 cycles, paired-end reads) using an input concentration of 1.6 pM with 1% PhiX.

Raw sequencing files from multiple corresponding Illumina sequencing lanes were merged and BCR sequence reads were processed using Cell Ranger version 3.1.0 (10x Genomics, with murine V(D)J reference GRCm38 (mm10)). Only cells with productive heavy and light chain sequences were retained, in case there was more than one productive heavy or light chain present per cell, the most abundant sequence (in terms of reads) was kept. Cells with IgG subtypes were used for overlap analysis and clones were defined based on V-, J-gene identity and identical combinations of CDRH3 + CDRL3 amino acid sequences.

### Generation and screening of monoclonal antibodies by yeast display

For the detection of antigen binding clones with known natural pairing of VH and VL chains, we used shared antibody sequences obtained by single-cell antibody repertoire sequencing. We expressed a subset of the 26 overlapping IgG+ antibody sequences that were at least present in three organs. Clonal grouping of the 26 IgG+ antibody sequences resulted in 17 clonal groups out of which at least one variant was chosen to be expressed as scFv. The selection criteria for choosing the sequence were: sequence that was shared with the majority of organs and displayed the highest abundance in bone marrow within the corresponding clonal group. Briefly, full length sequences were extracted from the bone marrow (or spleen) filtered_contig.csv files and aligned with IgBlast (*56*) before assembling full length scFvs containing VL and VH separated by a (Gly_4_Ser)_3_ linker and 30 bp homology overhangs on each side, which were ordered from Twist Bioscience. Gibson assembly for each scFv was performed using the NEBuilder HiFi DNA Assembly Master Mix (NEB, E2621L) according to manufacturer’s instructions with BamHI linearized pYD1 vector and scFv fragment in 1:4 ratio (in molarity). 5µl of each reaction including a negative control without scFv was subsequently transformed into 100 µl Fast Transformation of Mix & Go Competent Cells (Zymo T3001) each and plated on LB plates containing Ampicillin. After overnight incubation at 37°C, all plates, except the negative control, had >100 of colonies of which two were picked per plate to confirm correct insertion and sequence of scFvs via Sanger sequencing. After confirmation of successful scFv integration into pYD1-BamHI, plasmid DNA was extracted for each sample from a 1 ml overnight culture using the QIAprep Spin Miniprep Kit (Qiagen, 27104). Next, plasmid DNA was transformed into yeast cells using frozen-EZ Yeast Transformation II kit (Zymo T2001). Briefly, 0.2 µg of DNA was mixed with 25 µl of competent cells and 250 µl EZ3 solution. After a 45 minutes incubation at 37°C, cells were plated onto SD-CAA + 2 % glucose plates and incubated at 30°C for 3 days. Per plate, two colonies were inoculated in YNB-CAA + 2% glucose medium overnight at 30°C, followed by 24 hours induction in YNB-CAA + 2% galactose medium prior to flow cytometry screening. Staining of monoclonal scFv yeast cells was performed as described in the method section ‘Fluorescence activated cell sorting of scFv yeast libraries’. Briefly, 1x10^6^ cells were stained with μg/μL anti-FLAG-PE and 0.01 μg/μL RSV-F-AF647. Yeast cells expressing the RSV-F binding monoclonal antibody Palivizumab as scFv were used as positive control (AF647+/PE+) as well as negative control (PE+ only) to set the gates. Flow cytometry scanning to screen for double positive (AF647+/PE+) cells was performed on BD LSR Fortessa (BD Biosciences).

### Data visualization and statistical testing

Clonal expansion profiles (**Fig. 1D**) displaying the frequencies of clones and clonal fractions were generated using tcR R package (*65*). Clonal tracking plots (**Fig. 5D** and **fig. S9A**) were generated using R package immunarch (*50*). Heatmaps (**Fig. 1E, fig. 3C, Fig. S2B**, and **Fig. S4B**) were generated using R package pheatmap 1.0.12 (*66*), all other plots were generated using the R package ggplot2 3.3.2 (*67*), unless stated otherwise. The following R packages were used for visualization: RColorBrewer R package (*68*), cowplot (*69*). GraphPad Prism (version 7) was used for analysis of ELISA data and MAF subsampling analysis (**Fig. S1, A** and **F**) and for comparison between groups, t-test was used with p < 0.05 being considered as statistically significant. Violin plots and boxplots represented the median and interquartile range while other statistical data presented mean ± SEM.

## Acknowledgements

We acknowledge the ETH Zurich D-BSSE Animal Facility, in particular M.D. Hussherr and Dr. G. Camenisch for excellent assistance with animal experiments and with animal authorization. We acknowledge the ETH Zurich D-BSSE Genomics Facility Basel and Single Cell Unit, in particular E. Burcklen, I. Nissen, Dr. C. Beisel, Dr. M. Di Tacchio, and Dr. T. Horn for excellent support and assistance. We thank Mason Minot for scientific discussions.

## Author contributions

L.C. and S.T.R. designed experiments and overall study. L.C. performed mouse experiments, cell sorting and deep sequencing library preparations. L.C. and D.N. performed single-cell sequencing library preparations.

L.C. performed yeast display screenings. J.M.T and D.N. generated critical material and provided technical advice and support for yeast display experiments. L.C. and K.H. performed antibody repertoire analysis. S.F., C.R.W., and A.K. provided advice and support for repertoire analysis. F.S. expressed, purified and labelled RSV-F protein. L.C. and K.H. made figures. L.C., K.H., and S.T.R. wrote the manuscript. All authors reviewed and edited the manuscript.

## Data and materials availability

The raw FASTQ files from deep sequencing that support the findings of this study are deposited in the Sequence Read Archive (SRA) with the primary accession code PRJNA763201 (http://www.ncbi.nlm.nih.gov/bioproject/763201). Additional data and code that support the findings of this study are available on the GitHub repository: www.github.com/luciacsep/physiologicallandscape.

## Supplementary figures and tables

Figures (Fig. 1A, fig. 2 (legend), and fig. 5A and G) were created using BioRender.com.

**Fig. S1.**
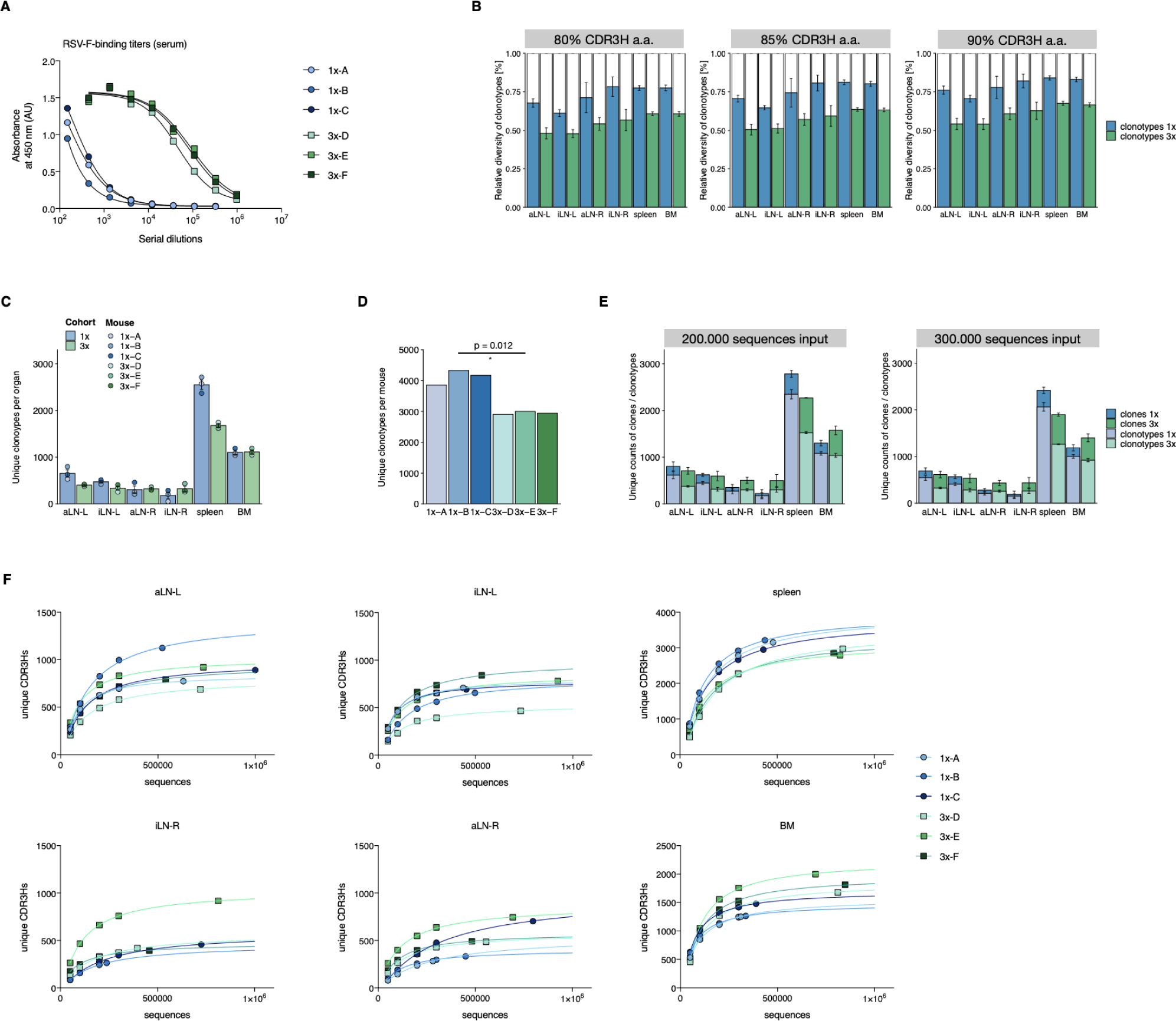
Serum titer measurements, evaluation of clonotyping, quantification of clonal diversities and subsampling analysis. (**A**) Serial dilutions of sera for the detection of RSV-F-specific titers ten days after single or final booster immunizations for cohort-1x and cohort-3x, respectively, using ELISA. A four-parameter logistic curve was fitted to the data by nonlinear regression. (**B**) Comparison of different clonotyping thresholds using 80%, 85%, and 90% CDRH3 a.a. similarities by plotting average diversities of unique clonotypes (numbers of unique clonotypes) relative to average diversities of unique clones (numbers of unique clones) for each organ per cohort (mean ± SEM, n=3). (**C**) Average clonotype diversity (mean ± SEM, n=3) by quantification of total numbers of unique clonotypes within each organ per mouse (point) and cohort (bar). (**D**) Clonotype diversity by quantification of total numbers of unique clonotypes across all organs within each mouse. For statistical analysis, Welch’s t-test was used on the average clonal diversity of cohort-1x mice and cohort-3x mice. (**E**) Comparison of varying sequencing read input (2x10^5^ and 3x10^5^) into the MAF pipeline by quantification of both average clonal and average clonotype diversities for each cohort (mean ± SEM). (**F**) Yield of unique clonal sequences (defined by identical CDRH3) after subsampling sequence input into the MAF pipeline (5x10^4^, 1x10^5^, 2x10^5^, 3x10^5^ and maximal sequence number per sample). Curves were fitted to the data by nonlinear regression. BM: bone marrow, aLN-L, -R: left and right axillary lymph nodes, iLN-L, -R: left and right inguinal lymph nodes.

**Fig. S2.**
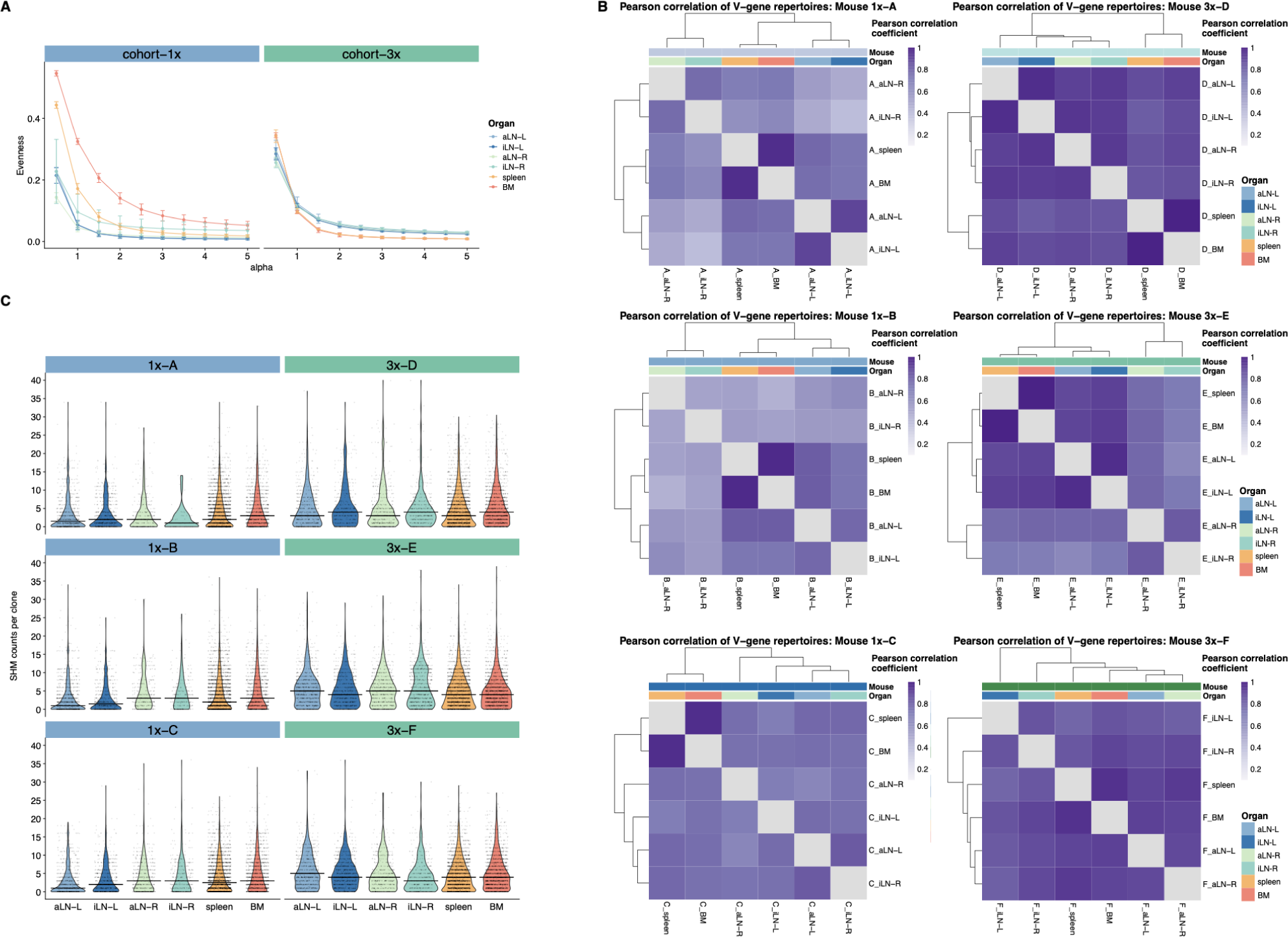
Average Evenness profiles, Pearson correlation of germline V-gene usage and quantification of somatic hypermutations. (**A**) To assess the distribution of clonal frequencies in the organ repertoires, we quantified clonal expansion profiles (Hill-based Evenness profiles) as previously reported (*1*). Average Hill-based Evenness profiles indicate the degree of clonal expansion of antibody repertoires within lymphoid organs of the two cohorts (mean ± SEM, n=3 per cohort) by using a continuum of diversity indices (the parameter alpha determines the impact of high frequency clones, with increasing alpha values, higher frequency clones are weighted more). For example, alpha = 1 indicates the Shannon Evenness, where higher values (up to 1) show uniform repertoires, whereas lower values (tending to 0) indicate polarized repertoires (*2*). Each colored line depicts the antibody repertoire of one organ (see legend). (**B**) Hierarchical clustering of pairwise Pearson correlations for the comparison of germline V-gene usage counts between organs for each mouse (**C**) Quantification of somatic hypermutation (nonsynonymous SHM counts per clone) for each organ repertoire of each mouse (horizontal line indicates the median, y-axis limit is set to 40). BM: bone marrow, aLN-L, -R: left and right axillary lymph nodes, iLN-L, -R: left and right inguinal lymph nodes.

**Fig. S3.**
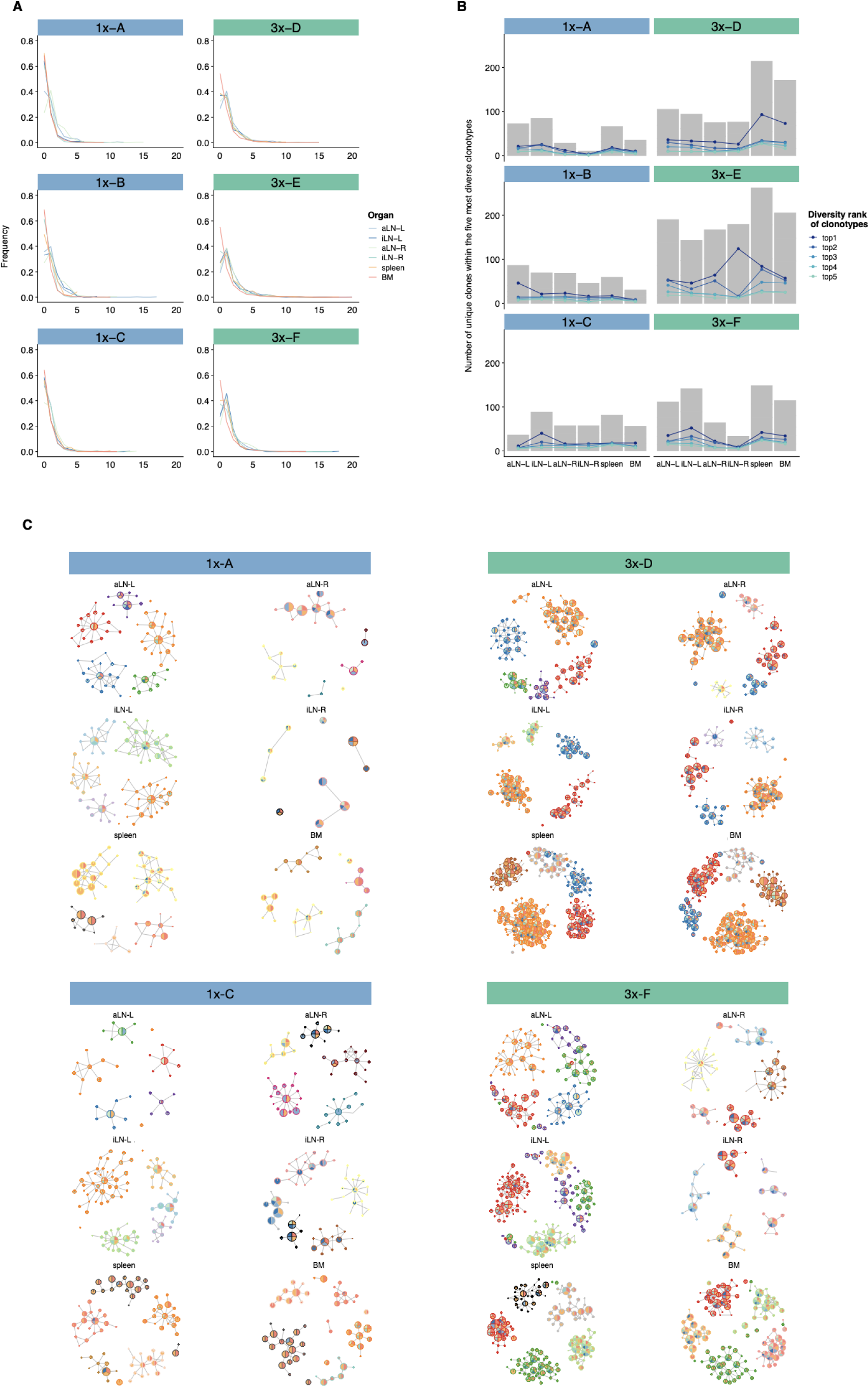
Quantification of clonal connectivity and clonal diversity and network-based analysis of the five most diverse clones. (**A**) Measurements of clonal connectivity degree distributions by quantifying the number of connections each node (clone) exhibits. Nodes belonging to the same clonotype with a Hamming distance of 1 were connected. Each line represents an organ. (**B**) Clonal diversities of the five most diverse clonotypes by quantifying the number of unique clones (clonal variants) within each clonotype in each organ. Each line represents one of the top 1-5 diversity ranked clones (see legend; the line indicates the diversity-rank of the top 1-5 most diverse clonotypes for each organ, it does not indicate the identical clonotype across organs). Grey bars represent the sum of all clonal variants within top 1-5 most diverse clonotypes for each organ. (**C**) Graphs show sequence-similarity networks of the five most diverse clonotypes within each lymphoid organ for mice 1x-A, 1x-C, 3x-D, and 3x-F. Each node represents a unique clone and all clones belonging to the same clonotype with a Hamming distance of 1 are connected (edges). Clones within each clonotype display the same border color throughout all organs within each mouse. The size of each clone represents the clonal frequency (in log) within the corresponding organ. Pie charts within each node depict organs that share the corresponding clone. BM: bone marrow, aLN-L, -R: left and right axillary lymph nodes, iLN-L, -R: left and right inguinal lymph nodes.

**Fig. S4.**
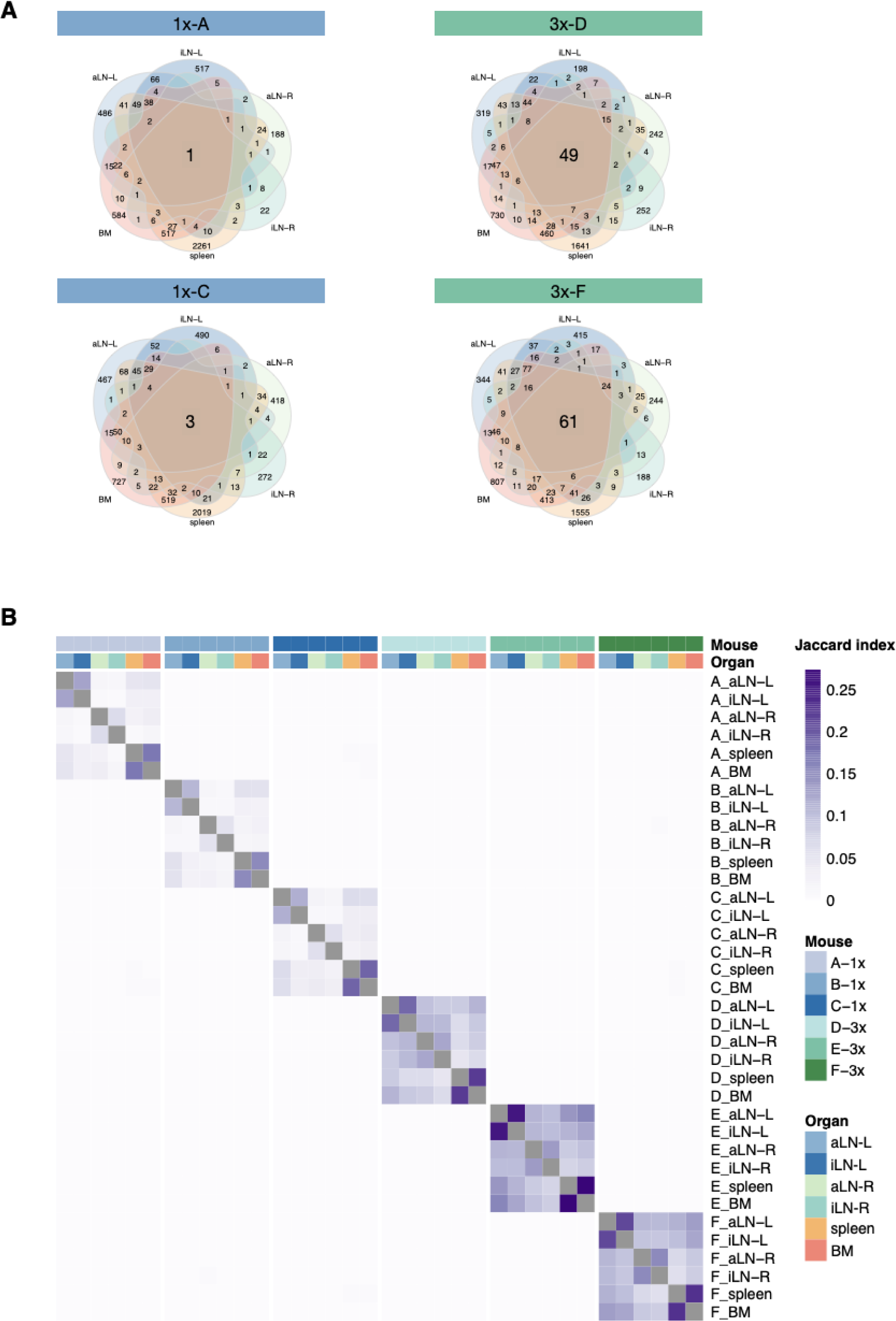
Repertoire similarity based on clonal overlap. (**A**) Venn diagrams depicting numbers of shared unique clones within mice 1x-A, 1x-C, 3x-D, and 3x-F. (**B**) Heatmap depicting pairwise Jaccard index to assess repertoire similarity based on overlap of unique clones across all samples. The pairwise Jaccard index is calculated by quantifying the size of intersection between two samples and dividing it by the length of the union of the same sample. BM: bone marrow, aLN-L, -R: left and right axillary lymph nodes, iLN-L, -R: left and right inguinal lymph nodes.

**Fig. S5.**
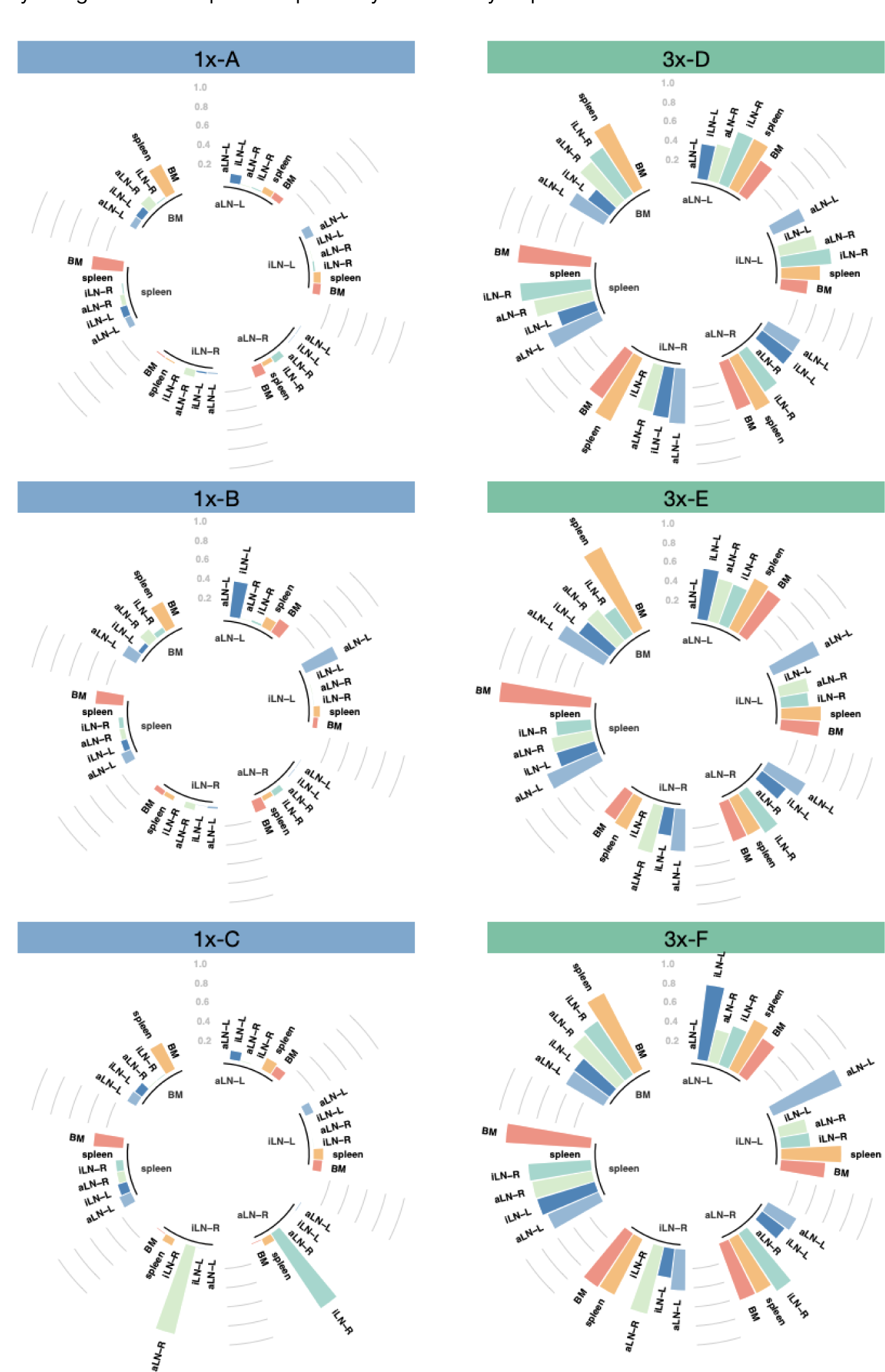
Repertoire similarity based on pairwise cosine similarities. Graphs depicting cosine similarities of clones between organ pairs for each mouse per cohort. Cosine similarity measurements represent the degree of repertoire similarity between two organs by including both clonal overlap and frequency information (by giving weight to clonal frequencies), ranging from 0 to 1. Each section of the plot represents one organ and each bar depicts the cosine similarity of the given organ to another organ of the same mouse. Higher cosine values correspond to higher repertoire similarities between two organs. BM: bone marrow, aLN-L, -R: left and right axillary lymph nodes, iLN-L, -R: left and right inguinal lymph nodes.

**Fig. S6.**
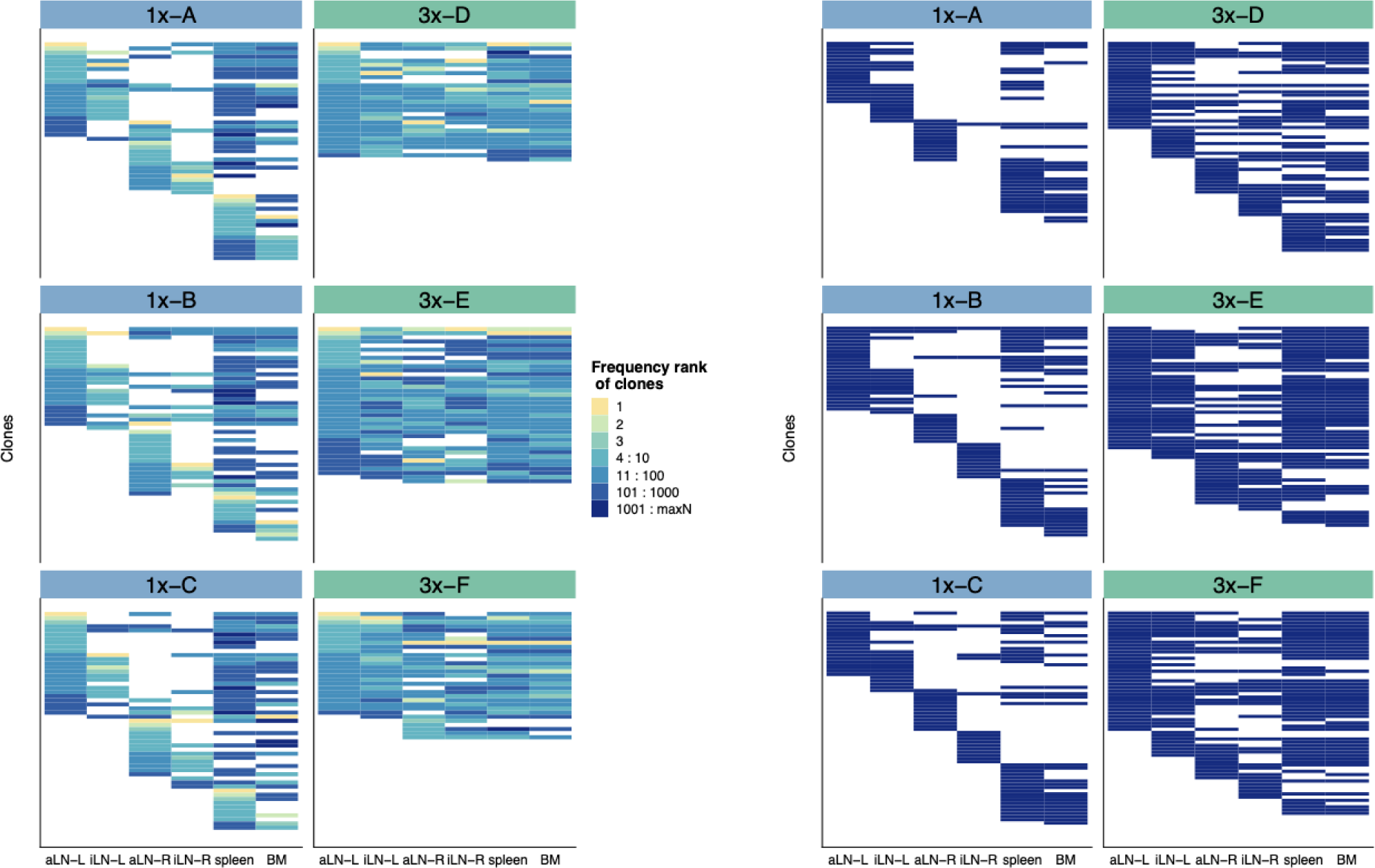
Tracking of diversity ranked clones across organs. (**A**) To dissect if clones that are shared between lymphoid organs display similar frequencies, we generated string plots (*3*). Tracking of the top-10 frequency ranked clones of each lymphoid organ across all six organs within each mouse is shown. String plots are based on Boolean values (presence or absence of a clone in the corresponding organ) with each column representing one lymphoid organ and each line representing one clone. Each line (clone) that is in the same position across columns (organs) represents an identical shared clone. The color encodes the frequency rank of the clone within the specific organ (see color legend). (**B**) Plots for tracking the top 100-110 frequency ranked clones of each lymphoid organ across all six organs within each mouse. BM: bone marrow, aLN-L, -R: left and right axillary lymph nodes, iLN-L, -R: left and right inguinal lymph nodes.

**Fig. S7.**
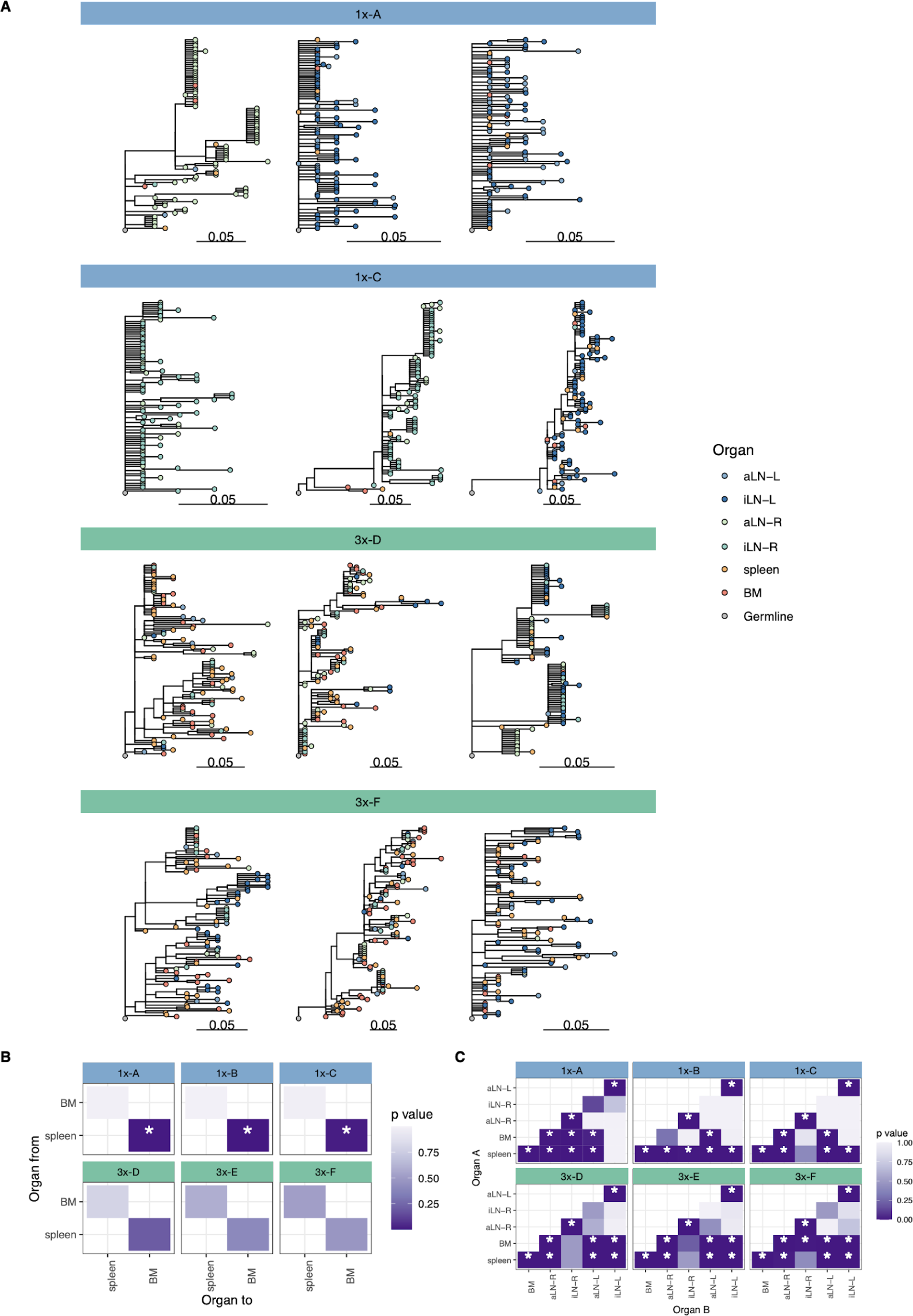
Phylogenetic analysis. (**A**) Phylogenetic trees of the three most diverse clonotypes from mice 1x-A, 1x-C, 3x-D, and 3x-F. Color of nodes indicates the lymphoid organ origin of each tip with grey nodes depicting the clonotype’s unmutated germline sequence. Scale bar shows somatic hypermutation per codon site. (**B**) Heatmaps displaying p-values of a switch proportion (SP) test, using only trees built from spleen and bone marrow sequences to quantify enrichments of transitions within clonotypes between spleen and bone marrow. (**C**) SP test permuting organ labels among trees and quantifying enrichment of transitions in either direction. BM: bone marrow, aLN-L, -R: left and right axillary lymph nodes, iLN-L, -R: left and right inguinal lymph nodes.

**Fig. S8.**
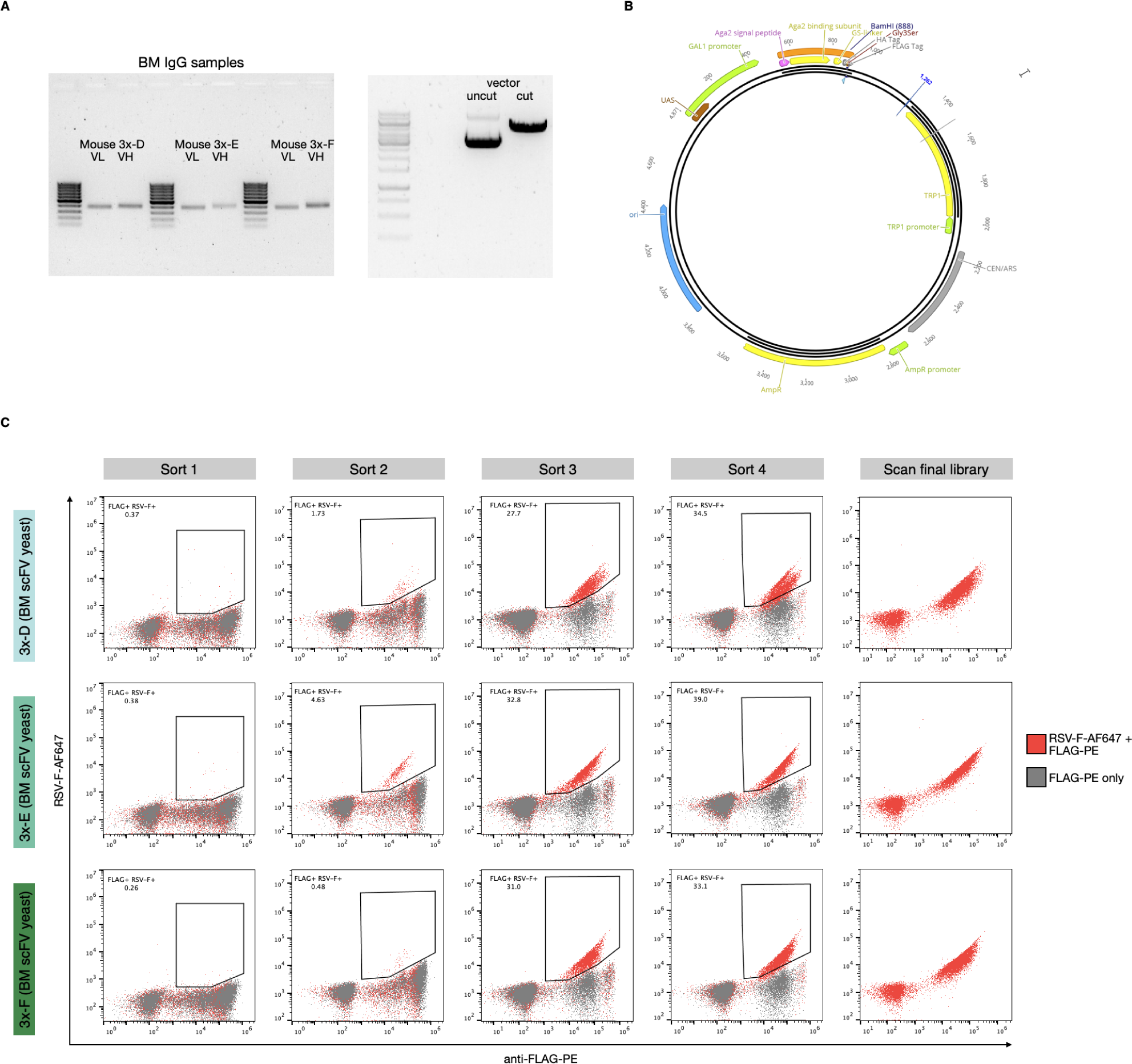
High-throughput antigen-specificity screening using scFv yeast display libraries. (**A**) Representative gel plots showing amplified VL and VH genes of bone marrow (BM) samples for cohort-3x mice (3x-D, 3x-E, 3x-F) as well as BamHI restricted yeast surface display vector. (**B**) Schematic of yeast surface display vector used for generation of scFv yeast display libraries. (**C**) FACS enrichment of RSV-F-specific binders using scFv yeast display. Representative flow cytometry dot plots show scFv yeast display library screenings for BM samples 3x-D, 3x-E, 3x-F by four progressive enrichment steps for FLAG-PE+ (x-axis) and RSV-F-AF647+ (y-axis) double positive cells (red, sort1 - sort4). Negative controls containing cells stained for FLAG expression only (grey) were used to set the sorting gates. Representative flow cytometry plots are shown to illustrate successful enrichments. The final sorted populations of yeast cells were screened to confirm purity before sequencing libraries were prepared.

**Fig. S9.**
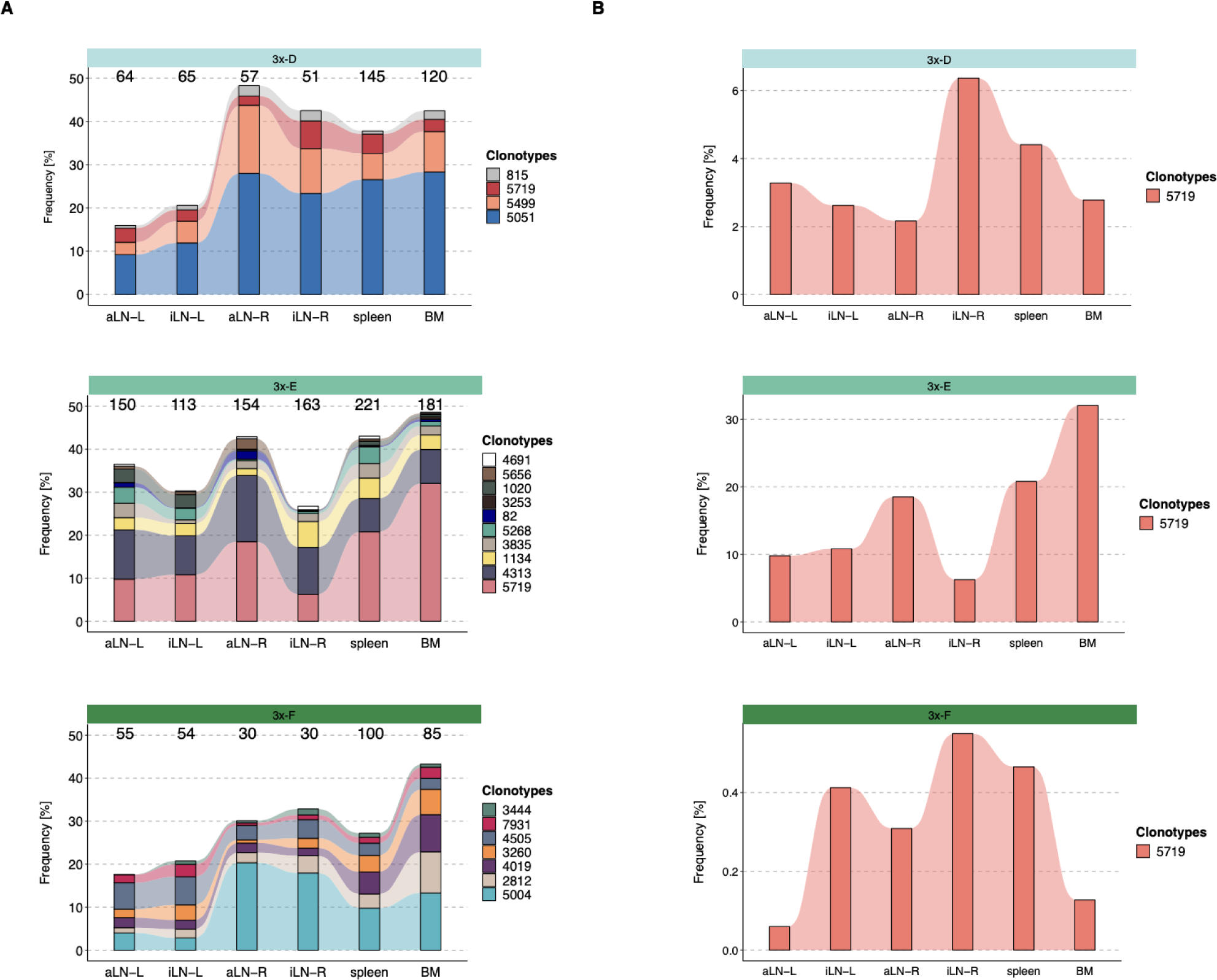
Tracking of clonotypes across all lymphoid organs. (**A**) Tracking of clonotypes that are associated with binding clones across all six lymphoid organs and their frequencies for each mouse. Numbers above each column indicate the numbers of unique clones belonging to the clonotypes within the corresponding organ (**B**) Frequency tracking of a public clonotype consisting of binding clones across all six lymphoid organs (y-axis display different ranges). BM: bone marrow, aLN-L, -R: left and right axillary lymph nodes, iLN-L, -R: left and right inguinal lymph nodes.

**Fig. S10.**
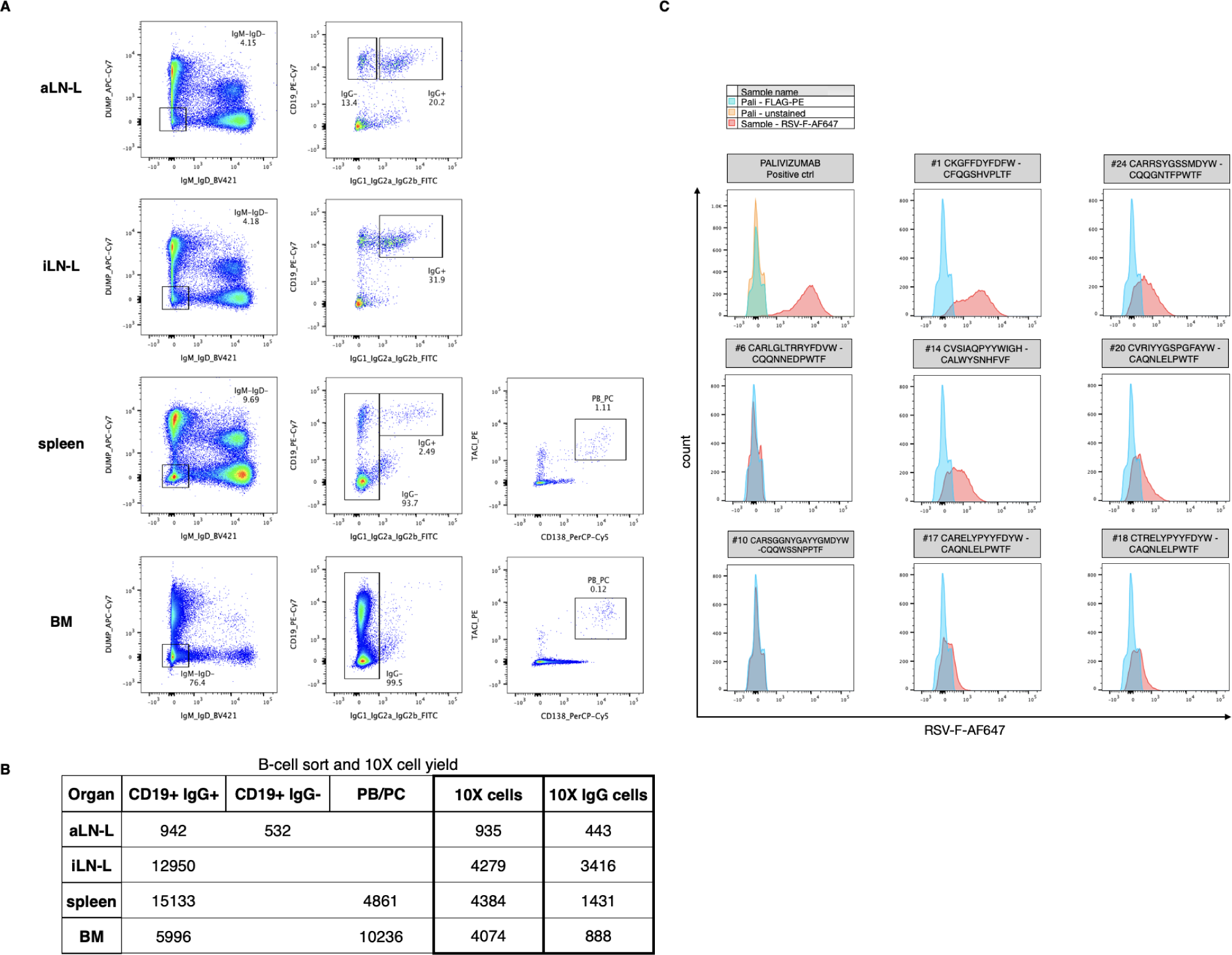
Isolation of B-cell subsets and screening of monoclonal yeast cells expressing scFvs. (**A**) Overview of the FACS strategy used for the isolation of specific B-cell subsets for each organ as input for single-cell sequencing. (**B**) Numbers of FACS isolated B-cell subsets per organ as well as yield of all cells and IgG+ B cells after single-cell V(D)J sequencing of antibody repertoires. (**C**) Flow cytometry histograms showing RSV-F-binding profiles of nine representative monoclonal single-cell scFv-expressing yeast clones, selected among the 18 tested clones according to overlap analysis (including a positive control expressing RSV-F-binding monoclonal antibody Palivizumab as scFv (see **table 1**)). Clones #1, 14, 17, 18, 20, and 24 were considered RSV-F-specific, whereas clones #6 and 10 are representatives for non-binding profiles. Clones were pre-gated on FLAG-PE expression. BM: bone marrow, aLN-L: left axillary lymph node, iLN-L: left inguinal lymph node.

**Table S1.** Sequence input into molecular amplification fingerprinting (MAF) pipeline. Numbers of quality-processed and length-trimmed sequencing reads as input into the MAF pipeline for error and bias correction for mice 1x-A, 1x-B, 1x-C, 3x-D, 3x-E, and 3x-F. BM: bone marrow, aLN-L, -R: left and right axillary lymph nodes, iLN-L, -R: left and right inguinal lymph nodes.

| Number sequence input into MAF pipeline |  |  |  |  |  |  |
| --- | --- | --- | --- | --- | --- | --- |
|  | 1x-A | 1x-B | 1x-C | 3x-D | 3x-E | 3x-F |
| aLN-L | 405000 | 405000 | 405000 | 405000 | 405000 | 405000 |
| iLN-L | 405000 | 405000 | 405000 | 405000 | 405000 | 405000 |
| aLN-R | 281114 | 405000 | 405000 | 405000 | 405000 | 405000 |
| iLN-R | 16206 | 238220 | 405000 | 396147 | 405000 | 405000 |
| spleen | 405000 | 405000 | 405000 | 405000 | 405000 | 405000 |
| BM | 311438 | 336122 | 389748 | 405000 | 405000 | 405000 |

**Table S2.** Clonal output after MAF processing. Numbers of unique clones (and clonotypes; defined by antibody sequences possessing identical germline V- and J-genes and 90% CDRH3 a.a. identity and identical length) obtained after MAF processing. BM: bone marrow, aLN-L, -R: left and right axillary lymph nodes, iLN-L, -R: left and right inguinal lymph nodes.

| Unique clones after MAF processing |  |  |  |  |  |  |
| --- | --- | --- | --- | --- | --- | --- |
|  | 1x-A | 1x-B | 1x-C | 3x-D | 3x-E | 3x-F |
| **aLN-L** | 740 (531) | 1055 (798) | 777 (629) | 630 (377) | 861 (407) | 760 (420) |
| **iLN-L** | 702 (480) | 615 (421) | 684 (513) | 419 (256) | 702 (348) | 794 (409) |
| **aLN-R** | 282 (250) | 323 (207) | 570 (461) | 457 (303) | 675 (361) | 481 (300) |
| **iLN-R** | 55 (50) | 263 (205) | 374 (289) | 419 (257) | 810 (435) | 389 (284) |
| **spleen** | 3025 (2573) | 3166 (2714) | 2916 (2370) | 2503 (1746) | 2475 (1618) | 2480 (1673) |
| **BM** | 1252 (1042) | 1264 (1080) | 1479 (1189) | 1520 (1049) | 1846 (1189) | 1665 (1099) |

**Table S3.** Overlap of the five most diverse clonotypes across lymphoid organs. Table indicates overlap of the five most diverse clonotypes (top1-top5) across lymphoid organs within each mouse. The numbers represent organs (1: aLN-L, 2: iLN-L, 3: iLN-R, 4: aLN-R, 5: spleen, 6: BM) that share the same clonotypes, being among the top five in all indicated organs. BM: bone marrow, aLN-L, -R: left and right axillary lymph nodes, iLN-L, -R: left and right inguinal lymph nodes.

| Five most diverse clonotypes per organ |  |  |  |  |  |  |
| --- | --- | --- | --- | --- | --- | --- |
|  |  | top1 | top2 | top3 | top4 | top5 |
| Mouse 1x-A | aLN-L (1) | 2 |  |  |  |  |
|  | iLN-L (2) |  | 2 |  |  |  |
|  | iLN-R (3) | 4 | 4,5,6 |  |  |  |
|  | aLN-R (4) | 3 | 3,5,6 |  |  |  |
|  | spleen (5) | 3,4,6 | 6 |  |  |  |
|  | BM (6) | 3,4,5 |  |  | 5 |  |
|  |  | top1 | top2 | top3 | top4 | top5 |
| Mouse 1x-B | aLN-L (1) |  |  |  | 2 | 2 |
|  | iLN-L (2) | 1 |  |  | 1 |  |
|  | iLN-R (3) | 4,6 |  | 4,5,6 | 5,6 |  |
|  | aLN-R (4) |  |  | 3,5,6 |  | 3,6 |
|  | spleen (5) | 6 |  | 3,4,6 | 3,6 |  |
|  | BM (6) | 3,4,5 | 3,5 |  | 3,4 | 5 |
|  |  | top1 | top2 | top3 | top4 | top5 |
| Mouse 1x-C | aLN-L (1) | 2,5,6 |  |  |  |  |
|  | iLN-L (2) | 2,5,6 |  |  |  |  |
|  | iLN-R (3) |  |  |  |  | 4 |
|  | aLN-R (4) |  |  |  | 3 |  |
|  | spleen (5) | 6 | 1,2,6 | 6 | 6 |  |
|  | BM (6) | 5 | 1,2,5 | 5 | 5 |  |
| Five most diverse clonotypes per organ |  |  |  |  |  |  |
|  |  | top1 | top2 | top3 | top4 | top5 |
| Mouse 3x-D | aLN-L (1) | all | all | all |  |  |
|  | iLN-L (2) | all | all | all |  |  |
|  | iLN-R (3) | all | all |  | all |  |
|  | aLN-R (4) | all | all |  |  | all |
|  | spleen (5) | all | 6 | all | 6 | all |
|  | BM (6) | all | all | 5 | 5 | all |
|  |  | top1 | top2 | top3 | top4 | top5 |
| Mouse 3x-E | aLN-L (1) | 2,5,6 | all | all | 2,4,5,6 |  |
|  | iLN-L (2) | all | all |  | 14,5,6 | 1,5,6 |
|  | iLN-R (3) | all | all |  |  |  |
|  | aLN-R (4) | all |  | all | 1,2,5,6 |  |
|  | spleen (5) | all | all | 1,2,6 | 1,2,4,6 |  |
|  | BM (6) | all | all | 1,2,5 | 1,2,4,5 |  |
|  |  | top1 | top2 | top3 | top4 | top5 |
| Mouse 3x-F | aLN-L (1) |  |  | all | 2,5,6 | 2 |
|  | iLN-L (2) | all | 5,6 | 1 | 1,5,6 | 3,6 |
|  | iLN-R (3) | 4 | 2,6 |  | 4,6 | all |
|  | aLN-R (4) |  |  | 3 | all | 3,6 |
|  | spleen (5) | 1,2,6 |  |  | all | 2,6 |
|  | BM (6) | 1,2,5 | all | 2,5 | 3,4 | 2,3 |

**Table S4.**
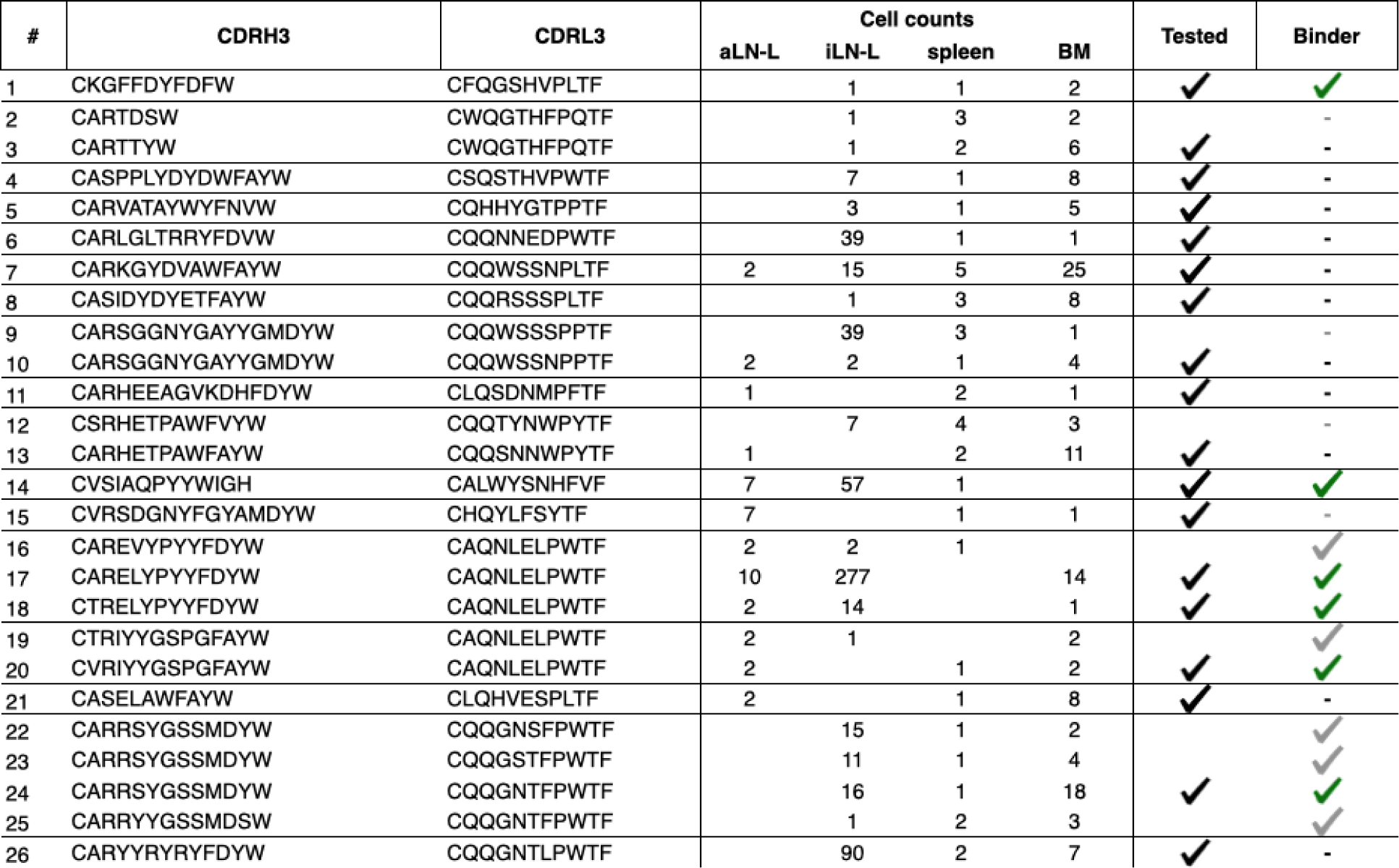
Overlap analysis for the identification of antigen-specific antibody sequences. CDRH3 and CDRL3 information of IgG+ B cells shared among at least three organs, including cell counts for each organ. Black check marks indicate antibodies tested for RSV-F-binding as scFv format in yeast cells. Green check marks indicate confirmed RSV-F-binding of single-cell clones, whereas grey symbols indicate presumptive binding / non-binding of clones belonging to the corresponding clonotype. BM: bone marrow, aLN-L: left axillary lymph node, iLN-L: left inguinal lymph node.

**Table S5.**
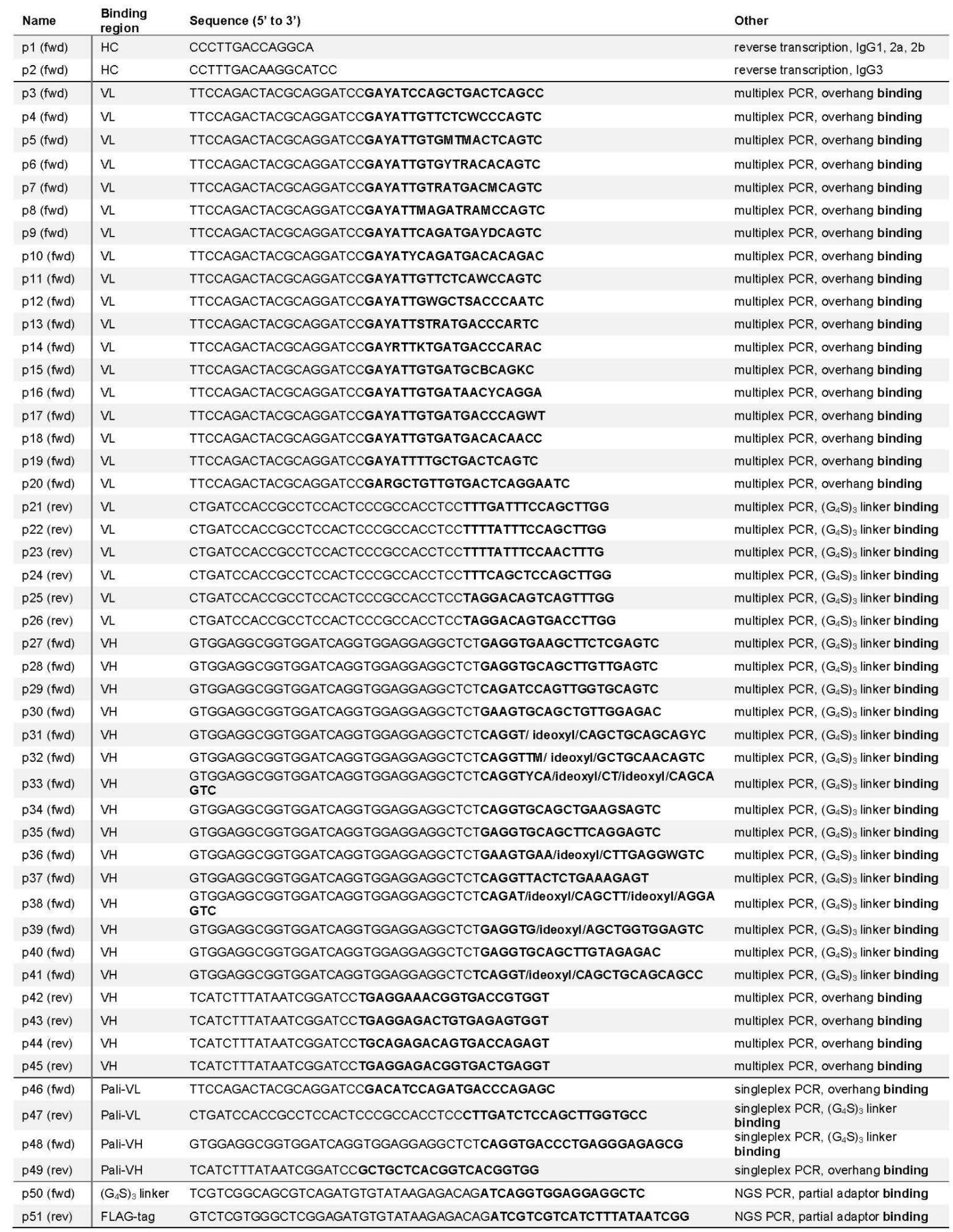
Primers used for yeast display screening and NGS library preparation. VL-binding primers p3-26 were adapted from Reddy et al., 2010 (*4*) and IgG- and VH-binding primers p1, 2, 27-45 were adapted from Khan et al., 2016 (*5*).

